# Evolution of the enhancer-rich regulatory region of the *POU1F1* gene

**DOI:** 10.1101/2023.07.26.550712

**Authors:** Michael Wallis, Qianlan Xu, Michal Krawczyk, Dorota Skowronska-Krawczyk

**Affiliations:** Department of Biochemistry and Biomedicine, School of Life Sciences, University of Sussex, Brighton BN1 9QG. UK; Department of Physiology and Biophysics, Department of Ophthalmology, Center for Translational Vision Research, School of Medicine, University of California, Irvine, CA, USA

**Keywords:** enhancers, super-enhancer, evolution, alternative splicing, promoter, comparative, pituitary

## Abstract

Precise spatio-temporal expression of genes in organogenesis is regulated by the coordinated interplay of DNA elements such as promoter and enhancers present in the regulatory region of a given locus. POU1F1 transcription factor plays a crucial role in the development of somatotrophs, lactotrophs and thyrotrophs in the anterior pituitary gland, and in maintaining high expression of growth hormone, prolactin and TSH. In mouse, expression of *POU1F1* is controlled by a region fenced by two CTCF sites, containing 5 upstream enhancer elements, designated E-A (5’ to 3’). We performed comparative sequence analysis of this regulatory region and discovered that three elements, B, C and E, are present in all vertebrate groups except Agnatha. One very long (>2kb) element (A) is unique to mammals suggesting a specific change in regulation of the gene in this group. Using DNA accessibility assay (ATAC-seq) we showed that conserved elements in anterior pituitary of four non-mammals are open, suggesting functionality as regulatory elements. We showed that, in many non-mammalian vertebrates, an additional upstream exon closely follows element E, leading to alternatively spliced transcripts. Here, element E functions as an alternative promoter, but in mammals this feature is lost, suggesting that conversion of alternative promoter to enhancer could be one evolutionary mechanism for enhancer birth. Our work shows that regulation of *POU1F1* changed markedly during the course of vertebrate evolution, use of a small number of enhancer elements combined with alternative promoters in non-mammalian vertebrates being replaced by use of a unique combination of enhancers in mammals.

## Introduction

The key DNA sequence involved in regulation of gene expression is the promoter, located immediately upstream of the transcription start site. To this region bind RNA polymerase and transcription factors (TFs) involved in regulating gene activation and transcription. Additional regulatory DNA elements such as enhancers, are involved in precise tissue-, cell type- and time-specific expression of the gene in eukaryotes. Enhancers are located upstream or downstream of their associated gene promoter and bind TFs which interact in turn with proteins bound to the promoter to execute their function. Enhancers are often located at a considerable distance from their target genes (many kb); their interaction with the promoter requires looping out of intervening DNA (1,2). In the case of some genes, several well-spaced enhancer elements appear to function in a coordinated manner, and together are referred to as a superenhancer (3). Regulation of gene expression may also be achieved by use of more than one promoter, associated with different starting exons alternatively spliced to the rest of the gene (4).

Despite their considerable functional differences, promoters and enhancers share many structural and functional traits (5). When activated, both regions display DNA accessibility, allowing different factors and co-factors to bind to their cognate sites. Both regions share several histone modifications (e.g. H3K4me1, H3K4me2, H3K27Ac), albeit at different levels. Moreover, both regions frequently bind the same factors, to exert their respective functions and both bind RNApolII which transcribes RNAs in both direction, although with significantly different efficiencies. While at the enhancer the levels of transcripts are low and rather equal in both directions, at the promoter, the coding transcript is transcribed with much higher efficiency than the transcript going in the reverse direction. Taken together, enhancers and promoters share many common architectural and functional features, which may provide clues about the evolutionary origin of enhancers.

POU1F1, also known as Pit1, is a POU-family TF which plays a key role in regulating development of the anterior pituitary gland, and the expression of growth hormone (GH), prolactin and thyrotropin (TSH) (6,7). While it is highly expressed in certain parts of the developing pituitary and mature cells producing these three hormones, its expression is virtually absent in most other tissues, though low levels in a few cell types may still be of significance. Extensive characterization of the promoter immediately upstream of *POU1F1* has been carried out in some mammals, confirming its crucial role in expression regulation (8). In addition, an element 10kb upstream of the promoter has been shown to be indispensable in maintaining high expression of *POU1F1* in adult mouse tissue. Detailed studies using transgenic animals have shown the importance of several upstream elements in spatio-temporal expression of *POU1F1* during pituitary development (9-11) including the long (>2kb) element located ∼6kb upstream of transcription start site (TSS), indispensable for early activation of the POU1F1 gene expression (12). This particular element, called EEα, is bound by ATBF1, a massive, multiple zinc-finger/homeodomain transcription factor, expressed specifically during early brain development. The identification of several discrete elements regulating the POU1F1 gene within a 25kb range from the TSS raises the possibility that these elements together comprise a superenhancer.

Multiple enhancers and/or alternative promoters allow for complex and subtle regulation of gene expression. In the case of *POU1F1* this includes precise regulation of the spatial and temporal expression of the gene during development, and the sustained high level of hormone expression maintained in mature pituitary somatotrophs, lactotrophs and thyrotrophs. To understand the evolution of the regulatory region of *POU1F1* gene, we performed a comparative study exploring the sequences of the upstream region of this gene across vertebrates. Our findings reveal that a conserved pattern of enhancers is used to regulate POU1F1 expression with specific modifications found in different clades. Notably, in birds and some other non-mammalian vertebrates our data show that alternative splicing of upstream exons appears to play an important role in regulating the *POU1F1* gene, while in placental mammals, a more-complex use of enhancers applies and alternative splicing is not employed.

## Results

### The *POU1F1* regulatory region in tetrapods

To visualize open regulatory elements in Pou1f1 gene in mouse pituitary we have performed ATAC-seq on nuclei isolated from adult (5 month-old) male mice. DNA accessibility studies revealed a cluster of 5 open regions upstream of the mouse *POU1F1* gene promoter, corresponding to five potential enhancers (elements E-A) (**Fig. 1**). Enhancer elements A, B and C have been characterized in detail previously and correspond to elements EEα, EEβ, and DE (8-13). Elements D and E are upstream of these. Highly conserved CTCF elements upstream of element E and in intron 5 flank the set of enhancers and coding region of the gene.

**Fig. 1.**
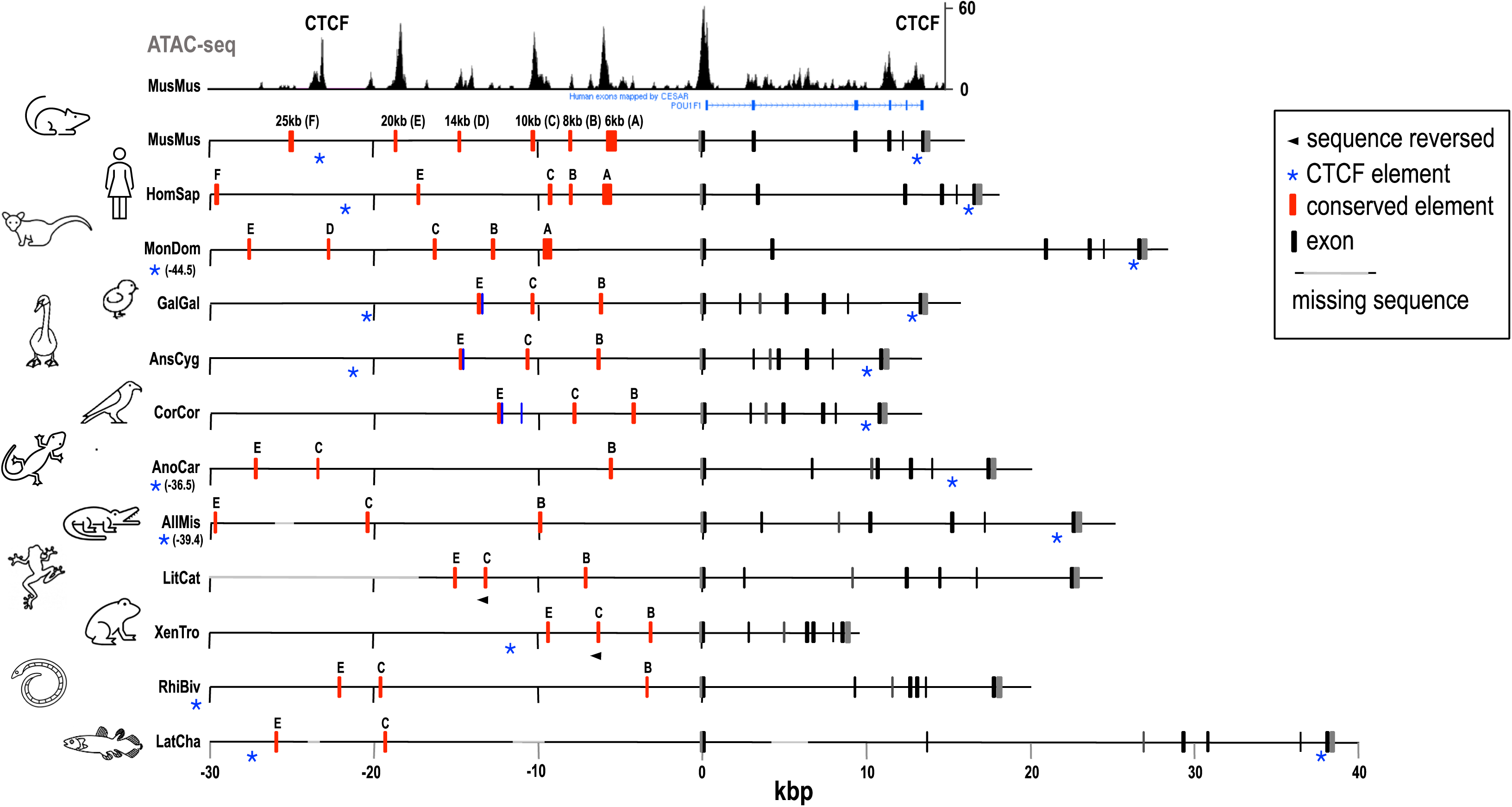
Organization of the *POU1F1* gene in representative tetrapods. Black and grey bars represent the 6 (mammals) or 7 (other vertebrates) exons. Red bars indicate conserved sequences (A-E) corresponding to the accessible regions in the ATAC profile (top panel). Blue bars represent alternatively spliced exon(s) found in birds (see text and Fig. 2); in mouse and human alternative upstream exon(s) are not used, but in the other species shown their usage or not is unknown due to lack of pituitary transcriptomes. F indicates a sequence upstream of the CTCF motif that is conserved in most placental mammals but not other tetrapods; it is not associated with an ATAC peak, and is probably not involved in *POU1F1* regulation. Sequences included are: MusMus, mouse, HomSap, human, MonDom, oppossum (mammals); GalGal, chicken, AnsCyg, swan goose, CorCor, crow (birds); AnoCar, anole lizard, AllMis, alligator (reptiles); LitCat, bullfrog, XenTro, clawed toad, RhiBiv, 2-lined caecilian (amphibia); LatCha, coelacanth (sarcopterygian fish). For full species names see *SI Appendix,* **Table S1**.

To assess how widely the type of complex regulatory region seen in mouse occurs, *POU1F1* genes in a wide range of tetrapod species were investigated, identifying conserved sequences by BLAST analysis. The five-enhancer arrangement was conserved in many mammals, including marsupials (**Fig.1**), but in a number of mammalian groups one, or sometimes two enhancers are lost. In reptiles, birds and amphibia, equivalents to only three of the five enhancers seen in mouse are conserved. Elements A and D are absent in all cases examined, but Elements B, C and E are present in almost all cases, suggesting a less flexible arrangement than seen in mammals (**Fig.1**). Some variation is apparent however. In particular, in Anura (frogs and toads) the orientation of element C is reversed compared with that in other amphibians and in all other tetrapods. Furthermore, the size of the regulatory region, and spacing of elements within it exhibited considerable variability. Most notably, in the axolotl, a salamander, the regulatory region extends over approximately 221kb, much greater than non-salamanders (cf 9-23kb for the amphibia included in **Fig.1**) and indeed other tetrapods examined. The introns of the axolotl *POU1F1* gene are also exceptionally large, though the spacing between promoter and exon 1 is similar to that in other amphibia. The large gene reflects the very large axolotl genome (14). Notably, the sequences of the enhancer elements and of coding sequence in axolotl are conserved and unremarkable (*SI Appendix,* **Fig. S1**) suggesting that the greatly increased gene size is not associated with functional changes. Finally, in human, enhancer element D is absent. A fuller assessment of the regulatory region across mammals is considered below.

### Alternative splicing and alternative promoters

A clear-cut transition in *POU1F1* regulation appears to have occurred during the course of evolution between mammals and birds, and potentially other non-mammalian tetrapods. In mammals transcription of *POU1F1* is controlled by up to 5 enhancers, comprising a potential superenhancer; there is no evidence for use of alternative promoters. Single cell studies (15) have shown that all 5 regulatory elements are accessible in each cell expressing the targets of POU1F1, further suggestive of cooperative work of all enhancers as a cluster in adult pituitary (*SI Appendix,* **Fig. S2**).

In non-mammalian tetrapods conserved sequences corresponding to just 3 of these enhancers are present. For a number of bird species pituitary transcriptomes are available. Analysis of these showed that in each case two alternative promoters are used, one equivalent to that seen in mammals (upstream of exon 1), the other equivalent to mammalian enhancer element E, upstream of an alternative exon, exon 0 (**Fig. 2**). These two promoters are used in approximately equal proportions. Exon 0 and exon 1 are spliced alternatively to exon 2. Analysis of a number of available mammalian pituitary transcriptomes (eutherians, including mouse, human and pig) confirmed that there is no alternative upstream exon - almost all transcripts start with exon 1. Reptiles and amphibia resemble birds in having just the three conserved upstream elements; whether they also use the alternative exon 0 could be established only for one amphibian for which a transcriptome is available, the salamander *Hynobius retardatus* (16). Analysis of this showed that exon 0 is present in this species with 24% exon 0-exon 2 splices and 76% exon1-ex2. A genomic sequence is not available for this species, precluding analysis of the sort shown in **Fig. 2**. For this species, as well as in birds, in-frame ATG codons would allow translation to give alternative POU1F1 proteins differing only in the short N-terminal sequences encoded by exons 0 or 1 (**Fig. 2C**).

**Fig. 2.**
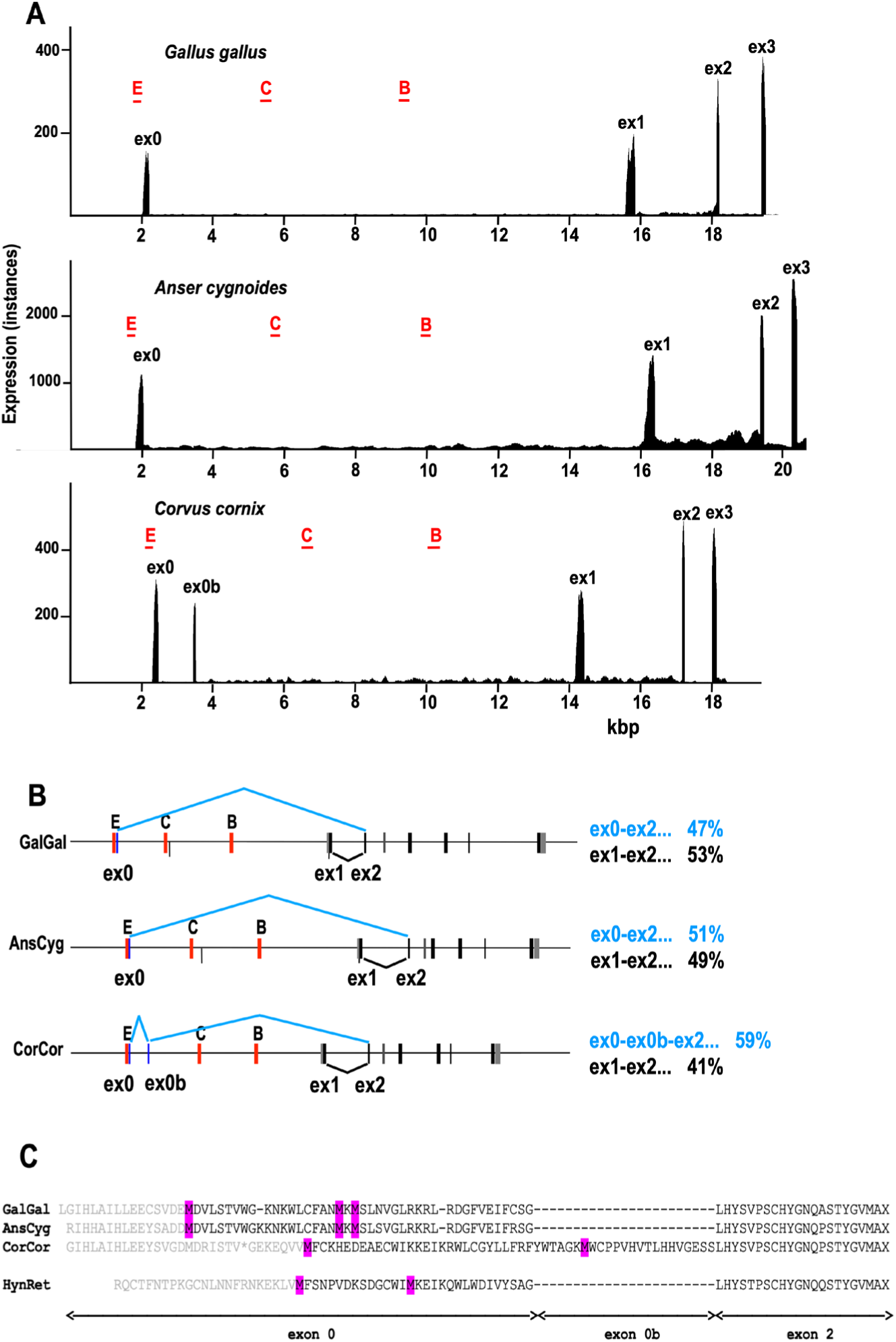
Alternative splicing at the 5’ end of the *POU1F1* gene in birds. **A**. Expression profiles for exons 1-3 and the upstream region in 3 bird species, illustrating the presence of an additional exon (ex0) in chicken (*Gallus gallus*) and swan goose (*Anser cygnoides*), and two additional exons (ex0 and ex0b) in hooded crow (*Corvus cornix*). E, C and B indicate the positions of the 3 conserved enhancer sequences, with enhancer E located immediately upstream of ex0 in each case. Based on analysis of SRA experiments SRX3216799 (chicken), SRX4891147 (swan goose), SRX330897 (hooded crow). **B**. Alternative splice patterns seen in the above 3 species. Percentages at the right are the proportions of the splice variants derived from the SRA experiments referred to above. **C**. Amino acid sequences (derived by conceptual translation) for exon 0 spliced to exon 2 for the 3 bird species indicated above and a salamander (*Hynobius retardatus)*. Potential translation start sites (methionine) are highlighted.

### *POU1F1* in fish

In teleost fish, identification of enhancer elements equivalent to those found in tetrapods initially proved difficult. However, elements B, C and E, but not A and D, were found in several other fish groups by Blast searches, and eventually were identified in many teleosts also. *POU1F1* was identified in all fish groups examined except Agnatha (lamprey and hagfish). Elements B, C and E were found upstream of this in most species examined, including lungfish, sturgeon, gar, many teleosts and all 10 Chondrichthyes represented in the NCBI wgs database (**Fig. 3**). Sequence similarity between tetrapods and fish is clear for each of elements B, C and E (*SI Appendix,* **Fig. S3**); in every case the order E-C-B (5’ to 3’) is retained, though relative spacing varies greatly. This last is most notable in the case of lungfish which has a very large genome (17) and a correspondingly large *POU1F1* gene (exon 1-exon 7 ∼541 kb) and upstream region (element E-exon 1 ∼523 kb; c.f. ∼5-38 kb in other fish). Element B was not found in coelacanth, probably due to a large unsequenced region between element C and exon 1.

**Fig. 3.**
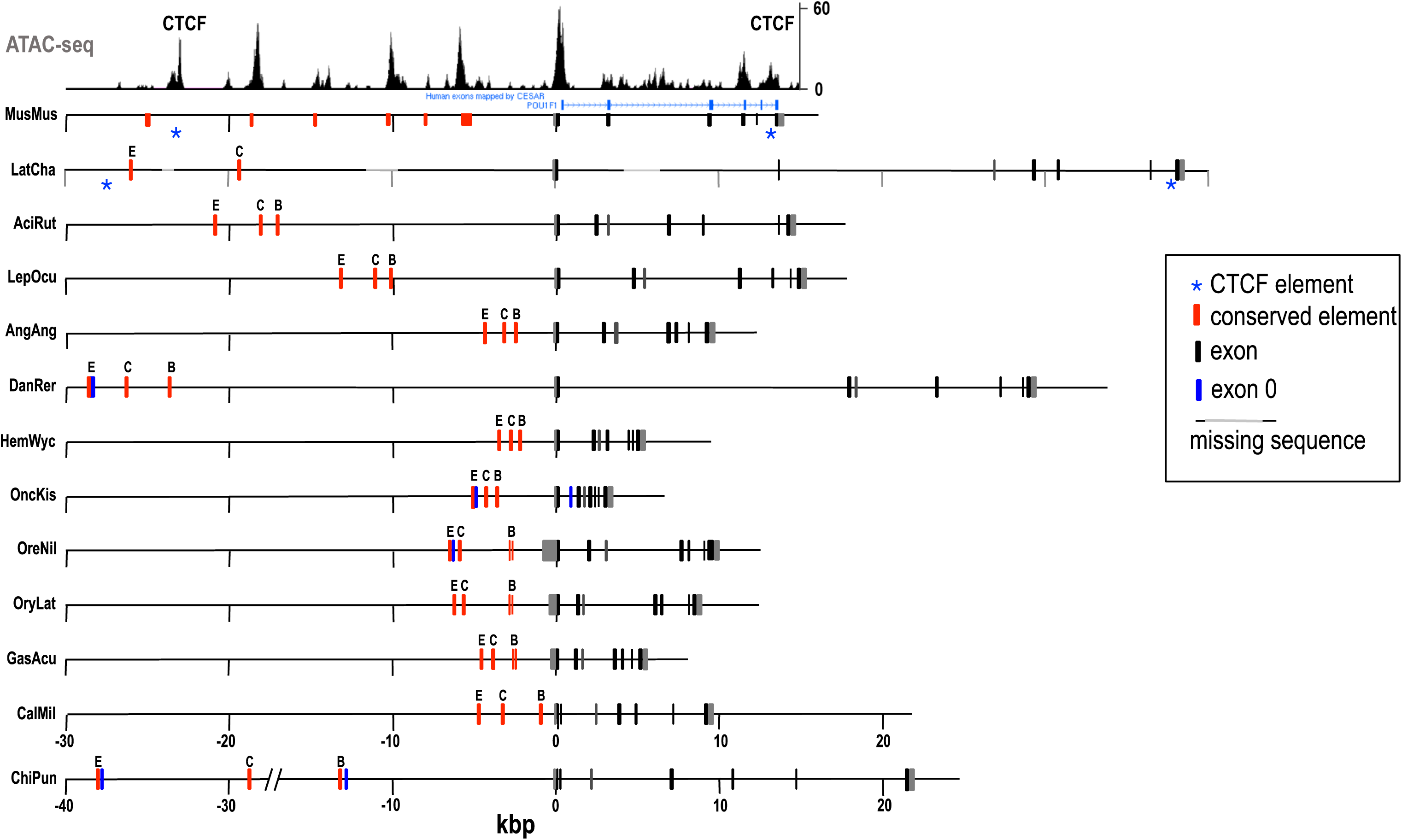
Organization of the *POU1F1* gene in fish. Black and grey bars represent the 7 exons. Red bars indicate conserved sequences (E,C,B) corresponding to the putative enhancers. Blue bars represent alternatively spliced upstream exon(s). In some cases where these are not shown (HemWyc, OruLat, GasAcu) analysis of transcriptomic data suggests that alternative upstream splicing does not occur, in others (LatCha, AciRut, LepOcu, AngAng, CalMil) presence or absence of upstream splicing cannot be evaluated owing to lack of appropriate transcriptomic data. Sequences included are: MusMus, mouse, LatCha, coelacanth, AciRut, sturgeon, LepOcu, spotted gar, AngAng, eel, DanRer, zebrafish, HemWyc, catfish, OncKis, salmon, OreNil, tilapia, OryLat, medaka, GasAcu, stickleback, CalMil, elephant shark, ChiPun, bamboo shark. For full species names see *SI Appendix,* Table S1.

In some fish *POU1F1* contains an additional, untranslated exon (exon 0) closely downstream of enhancer element E, similar to that seen in birds. In zebrafish and some close relatives, exon 0 is found in almost all transcripts; in others, for example salmon, it is found in a relatively small proportion (e.g. ∼6% in coho salmon, *Oncorhynchus kisutch*), with most transcripts initiating in exon 1.

### POU1F1 in mammals

Analysis of the mouse *pou1f1* gene showed the presence of 5 upstream enhancer elements, but the corresponding region of human revealed only four, with element D missing (**Fig. 1**). To assess the extent of such variability in mammals we examined the *POU1F1* gene in a wide variety of mammals, including almost all those for which genome sequences are available in the wgs database (∼300 species). In all of these a single *POU1F1* gene was identified, although in a few cases the sequence was incomplete. The 5-enhancer arrangement was found in at least some representatives of each of the main mammalian groups (Marsupialia, Afrotheria, Xenarthra, Laurasiatheria and Euarchontoglires) except Monotremata (**Fig. 4**; *SI Appendix,* **Fig. S4**). Enhancer elements A and C were detected in all mammals examined. Element B was present in all except elephant shrew (*Elephantulus edwardii;* Afrotheria) and gundi (*Ctenodactylus gundi*; Rodentia). On the other hand, elements D and E were absent from a number of groups. The organization of the regulatory region in a selection of mammals is shown in **Fig. 4**.

**Fig. 4.**
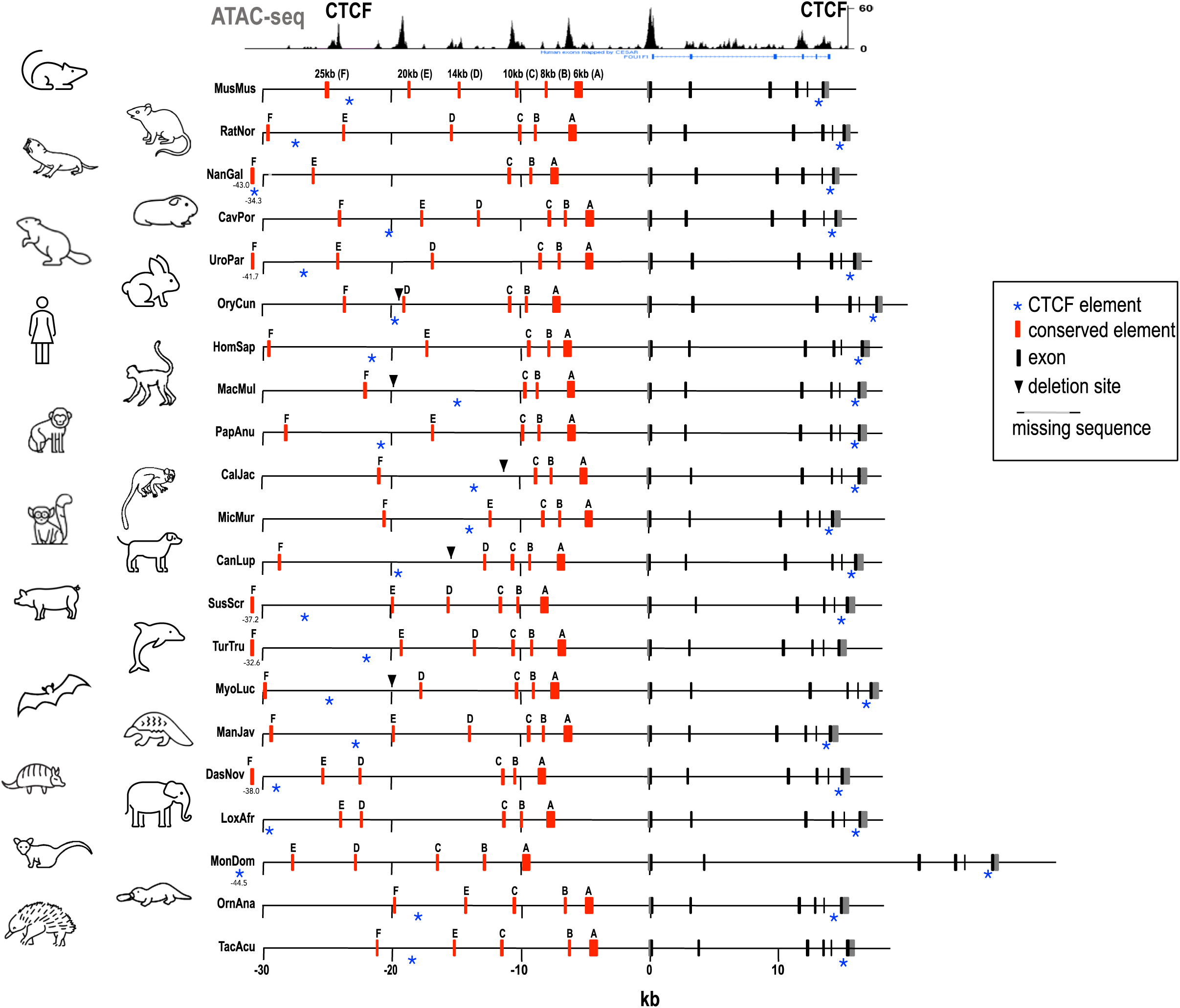
Organization of the *POU1F1* gene in mammals. Black bars represent the 6 exons. Red bars indicate conserved sequences (A-E) corresponding to the accessible regions in the ATAC profile (top panel). A vertical arrow indicates site of a deletion removing the E element. Sequences included are: MusMus, mouse, RatNor, rat, NanGal, blind mole rat, CavPor, guinea pig, UroPar, arctic ground squirrel, OryCun, rabbit, HomSap, human, MacMul, macaque, PapAnu, baboon, CalJac, marmoset, MicMur, mouse lemur, CanLup, dog, SusScr, pig, TurTru, dolphin, MyoLuc, little brown bat, ManJav, pangolin, DasNov, armadillo, LoxAfr, elephant, MonDom, oppossum, OrnAna, platypus, TacAcu, echidna. For full species names see *SI Appendix,* Table S1.

Element D is not present in the sequence upstream of *POU1F1* in monotremes (platypus and echidna). Since Monotremata is the outgroup for all other mammals this absence could be because monotremes represent an intermediate between the 3-element arrangement seen in most non-mammalian vertebrates and the 5-element structure of many mammals. On the other hand it is possible that the 5-element organization appeared early in mammalian evolution, and that element D was subsequently lost in Monotremata. Notably, the distance between elements C and E is considerably less in monotremes (∼3.8kb) than in most other mammals (e.g mouse, 10kb). Element D is also absent in all Primates and the two orders most closely related to primates, Scandentia and Dermoptera. This is clearly a consequence of loss of this element on this branch of Euarchontoglires, after separation from Glires (rodents and lagomorphs). Whether this was due to loss of element D by deletion or sequence divergence is not clear from the sequence data, given that it occurred at a fairly early stage of eutherian evolution. Element D was also absent in elephant shrew and two species in the rodent family Heteromyidae, pocket mouse (*Perognathus*) and kangaroo rat (*Dipodomys*).

Element E is present and well conserved in monotremes and marsupials. Indeed, in some marsupials, but not *Monodelphis* (opossum), it is duplicated. However, sequence conservation of Element E is relatively poor in Eutheria, and the element is missing in a number of groups. In most cases, this is clearly due to deletion. Thus in primates Element E is absent in all New World Monkeys (NWM) and in the group of Old World Monkeys (OWM) including *Macaca* and *Colobus*. Analysis of sequences shows that this is due to distinct, independent deletions. Independent deletions also removed element E in some rodents (Gliridae - dormice), all lagomorphs (rabbits and hares), some Carnivora (Canidae - dog family; sequence divergence high in other Carnivora), all Chiroptera (bats), several Eulipotyphla (moles and hedgehogs) and one Afrotherian (aardvark, *Orycteropus*). The lowered sequence conservation of element E suggests decreased importance in Eutheria, and the repeated deletions suggest that it may have been deleterious in some groups. Notably, deletion of element E in NWM and some OWM, in addition to the loss of element D in all primates, means that these groups retain only 3 enhancer elements, C, B and A. The hedgehog, *Erinaceus*, also retains only these 3 elements, while elephant shrew only has elements A, C and E.

### CTCF

A conserved CTCF binding site (18) was found upstream of the 5’-most enhancer element (element E except for those mammals in which this element is not present) in all tetrapods and in coelacanth and lungfish. This site is strongly conserved in Eutheria, but not monotremes or marsupials. An equivalent CTCF binding sequence was not found consistently in other fish. A conserved CTCF binding site is also found in the 3’-most intron in mammals, reptiles, birds, coelacanth and lungfish (**Figs. 1 and 4**). Alignments are shown in *SI Appendix,* **Figs. S1 and S4**.

### Promoter

The promoter region upstream of exon 1 of *POU1F1* has been characterized previously in rat (7,8) and shown to contain binding sites for POU1F1 itself, upstream (stimulatory) and downstream (inhibitory) of the transcription start site (8,19). The POU1F1 sites are strongly conserved in mammals and indeed in all tetrapods, but less so fish (*SI Appendix* **Figs. S1, S3 and S4**). The mouse promoter contains a TATA box sequence, which is strongly conserved, with the extended sequence TATAAATAC being found in most mammals, all birds and amphibia, all non-teleost fish and most teleosts (*SI Appendix,* **Figs. S1, S3-S5**). Minor substitutions, unlikely to affect function occur in reptiles and some mammals and teleosts. Thus in dog (TATAAGCGC) and pig (CATAATAC) *POU1F1* expression is as high as in mouse, rat and human (all TATAAATAC) (*SI Appendix,* **Table S2**). Significant substitutions, likely to abrogate function are seen in a few teleosts, including zebrafish, where obligate use of upstream exon 0 would imply that the promoter in front of exon 1 is less or no longer functional.

### Expression levels

To determine whether changes seen in the organization of the *POU1F1* upstream regulatory region during vertebrate evolution were associated with changes in overall expression of the gene, transcription in the anterior pituitary was assessed for a variety of species using transcriptomic data available from the NCBI Sequence Read Archive (SRA). Such data uses material from a wide range of studies, with animals of either sex and varying physiological states and subject to various experimental conditions. Results are shown in *SI Appendix,* **Table S2**. In all cases studied, expression levels were high, and there was no clear evidence for gross changes in expression level associated with varying organization of the regulatory region of the gene. This does not rule out more-subtle changes, for example reflecting physiological or developmental state.

### DNA Accessibility in non-mammals

Identification of additional potential regulatory elements upstream of *POU1F1* was initially based on the DNA accessibility profile in mouse (**Fig. 1**). The observation that accessibility is associated with sequence conservation was then used to identify sequence regions potentially involved in gene regulation in other species. In order to support this involvement, DNA accessibility studies were carried out on several non-mammalian species. Using the established protocol of ATAC-seq (20,21) we have assessed DNA accessibility in guinea fowl, alligator and bullfrog. In brief, we have dissected pituitaries of each animal, isolated nuclei and performed ATAC-seq according to the well established laboratory protocol (21,22). Two animal per group were analyzed. ATAC-seq raw reads data were mapped to specific genome and visualized using IGV software (**Fig. 5A**) (23,24). Data were presented together with schematic representation of the region for each animal and conserved elements. Our analysis demonstrated that the conserved regions in these species correspond to regions of high DNA accessibility in pituitaries, suggestive of functionally active regulatory elements.

**Fig. 5.**
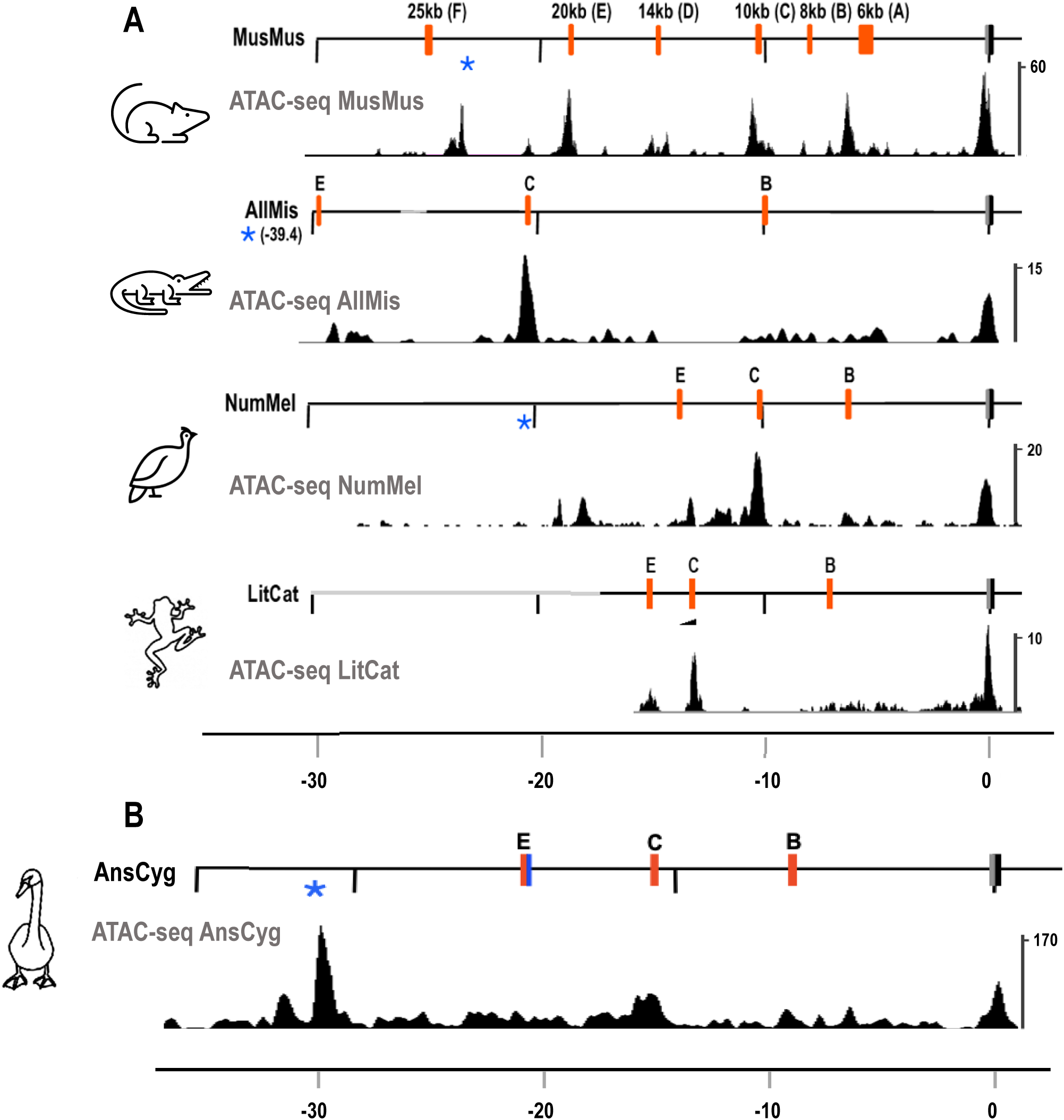
DNA accessibility upstream of the *POU1F1* gene in selected non-mammalian species, studied using ATAC. **A**. Examples of DNA accessibility in three species as compared to the mouse regulatory region (*top*). Alignment of open regions in alligator (AllMis), guinea fowl (NumMel) and bullfrog (LitCat) shows accessible DNA in regions corresponding to the conserved elements as found by the sequence similarities. B. DNA accessibility in the goose (Anser cygnoides, AnsCyg) (25) regulatory region showing conservation of the regions and their DNA availability for regulation.

Similarly, we used recently published goose pituitary ATAC-seq data (25) to align the conserved sequences with the DNA accessibility of goose *POU1F1* regulatory region. As in the case of guinea fowl, alligator and bullfrog, highly conserved regulatory elements were open in the goose pituitary, strongly suggesting the functionality of these regions (**Fig. 5B**).

### Enhancer alignments

Alignments for the five enhancer elements are shown in *SI Appendix,* **Figs. S1, S3 and S4**. In most cases, there is ‘structure’ within the alignment, with regions of relatively high similarity separated by poorly conserved regions. This is particularly marked in the case of element B where 4 such subregions can be seen. In some fish, including medaka and stickleback, these are more widely dispersed, and less distinct. Notably this structuring and spacing is retained in axolotl and lungfish, despite the drastically expanded genome in these species, with distances between enhancer elements increased almost 10-fold compared with other species.

Autoregulatory POU1F1 sites has been previously identified in the *POU1F1* promoter and enhancer element C in rodents (8,11). Many of these sites are strongly conserved across tetrapods (*SI Appendix,* **Figs. S1, S5**). Thus, of the 5 POU1F1 binding sites identified on enhancer DE in mouse (12), three are identifiable and conserved in the element C alignment (*SI Appendix,* **Figs. S1, S5**), but the other two are upstream of this in poorly conserved sequence. The two POU1F1 sites identified in the promoter (8) are well conserved. Potential POU1F1 binding sites were identified in element E, where they are strongly conserved in non-mammalian tetrapods, but not in placental mammals (*SI Appendix,* **Fig. S5**). The mammalian-specific elements A and D also contain potential POU1F1 binding sites, with up to three in element A, though these are poorly conserved (*SI Appendix,* **Fig. S5**). The presence of strongly conserved POU1F1 binding sites in element E in non-mammals that are much less well conserved in placental mammals, suggests that autoregulation of this element by POU1F1 may be important while it functions as a promoter, but less so when this role is lost. Appearance of POU1F1 binding sites in elements D and A, unique to mammals, would allow the overall autoregulatory role to be maintained and perhaps strengthened in mammals.

## Discussion

### Regulatory elements of the POU1F1 gene

Previous studies have established that expression of the *POU1F1* gene in mouse and rat is controlled by a number of transcription factors, including POU1F1 itself, via interactions with its promoter and at least three upstream enhancer sequences (6-13). Here we propose that in mouse there are 5 potential regulatory elements (enhancers) upstream of the mouse *POU1F1* promoter, on the basis of DNA accessibility and sequence conservation. These are designated elements E-A (5’ to 3’). They are flanked by CTCF binding sites upstream of element E and in intron 5 of the *POU1F1* gene, suggesting that they may constitute a cluster of enhancers cooperatively regulating expression of the locus (18). Element C corresponds to the previously-described definitive enhancer (DE; 11) while elements A and B correspond to the early enhancers (EEα, EEβ, respectively; 10,12,13).

In the study reported here, the nature of the regulatory elements associated with *POU1F1* was explored by examining genome sequences across vertebrate groups. The *POU1F1* gene was identified in all groups except the Agnatha (lamprey and hagfish). In all these groups, conserved sequences corresponding to enhancer elements B, C and E were detected, but elements A and D were only detected in mammals (Fig. 6). Notably, in many non-mammalian vertebrates for which pituitary transcriptomes are available (birds, salamander, teleosts and chondrichthyes) transcription can start at an upstream site, with use of an alternate exon (exon 0) which is spliced into exon 2 (birds, salamander, Fig. 2) or exon 1 (fish). In most cases transcripts starting at exon 0 are no more abundant than those starting at exon 1 (and in some fish, transcription from exon 0 was not apparent), but in zebrafish exon 0 appears to be used exclusively. In all cases exon 0 lies closely downstream of element E, suggesting that the latter acts here as an alternative promoter. DNA accessibility studies supported the existence of the regulatory elements B, C and E in a representative bird, reptile and amphibian. There was no suggestion of use of exon 0 or another alternative upstream exon in placental mammals for which pituitary transcriptomes are available (several species of rodents, primates, carnivores and artiodactyls).

**Fig. 6.**
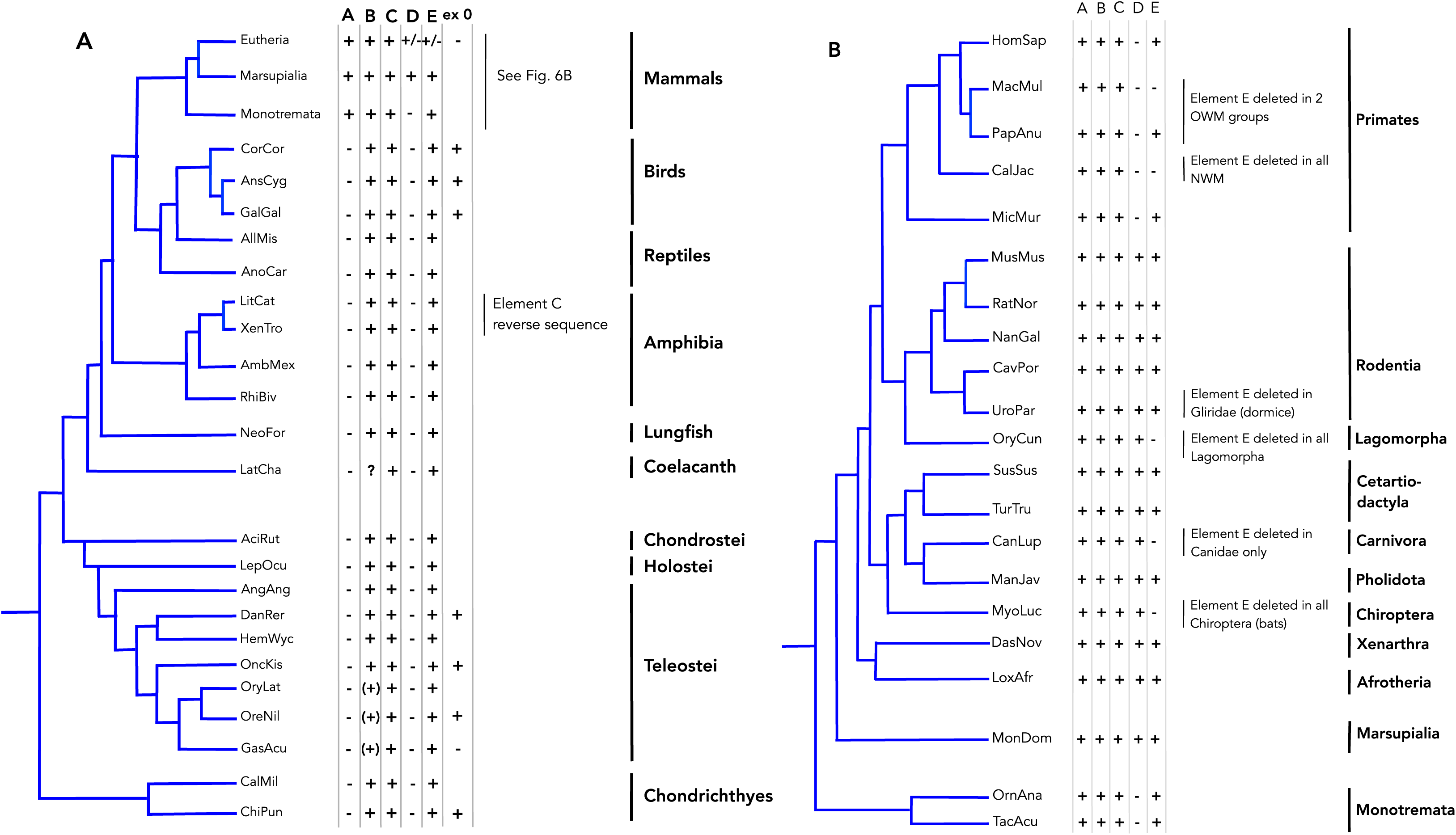
Phylogenetic trees summarizing the main events in *POU1F1* regulatory region evolution in (A) Vertebrates and (B) Mammals. Phylogenies are based on (35,36). Presence or absence of regulatory elements A-E in selected species is indicated.

### Evolution of the regulatory region

These observations suggest that the regulatory mechanisms for the *POU1F1* gene have been maintained fairly constant across all non-mammalian vertebrates except Agnatha, with use of alternative promoters playing an important role (**Fig. 6A**). With the emergence of mammals substantial changes occurred, with the acquisition of new regulatory elements (elements A and D) and loss of the alternative exon associated with element E, the latter potentially acquiring function as an enhancer. Element A must have been acquired early - it is found in all mammalian groups, but element D is not present in monotremes, so could have been acquired later.

The exact time in evolution of when exon 0 was lost is less clear. While, it is present in all bird species examined, sequence similarity (based on Blast analysis) suggests that it may also be present in reptiles, amphibia, monotremes and marsupials. Its presence in amphibia is supported by the pituitary transcriptome data of a salamander. Across mammals, the composition of the *POU1F1* regulatory region appears to be less stable/conserved than is seen for other vertebrates. Elements A, B and C are found in nearly all mammalian groups, but element D is absent in primates and the two groups most closely related to primates (tree shrews and flying lemurs), while element E is absent from a number of groups, reflecting at least 6 independent deletions (**Fig. 6B**). This suggests that element E, having lost its role as a promoter for exon 0, may play a rather equivocal role in mammals; where present its sequence is usually conserved, suggesting the retention of specific function. However, the fact that it has been completely removed by deletion on multiple separate occasions suggests that its presence may even have been disadvantageous in some cases. It is notable that in at least two primate groups (NWM and some OWM, including macaques) the deletion of element E and the absence of element D, results in the retention of only elements A, B and C, with no apparent effect on level of *POU1F1* expression. Notably, the variability of the regulatory region of the *POU1F1* gene in mammals contrasts with the strongly conserved nature of the coding sequence (26).

The most important change occurring during the evolution of *POU1F1* regulation appears to be that associated with the emergence of mammals. This involves loss of exon 0 and its associated alternative promoter, as well as acquisition of two additional enhancer elements, A and D. However, a number of additional features are also worth noting, including the switch in orientation of element C seen in Anura (frogs and toads) and retention of an apparently normal 3-element organisation in axolotl and lungfish, despite (independent) expansions of genome size with a ten-fold increase in distance between element E and promoter.

The origins of the 5 enhancer elements remain unclear. Elements B, C and E are clearly present in Chondrichthyes, and are well conserved across almost all vertebrate groups but, like the *POU1F1* gene, could not be detected in Agnatha. Sequence comparisons of the enhancer elements detected no similarity between them. The origin of the additional elements (A and D) seen in mammals is not clear. Element D is not present in monotremes; it may have appeared later in mammalian evolution or may have been present in the ancestor of monotremes and theria (placentals and marsupials), and subsequently lost in the former. The most interesting case is that of element E. Our data show that in birds and some amphibia and fish element E is immediately upstream of the additional 5’ exon (exon 0). It is possible that element E was originally an upstream enhancer that was taken over as a promoter in some groups. It is also possible that transcription from exon 0, under the control of the alternative promoter (Element E), allows separate regulation of *POU1F1* expression in early development in non-mammalian vertebrates, this function being taken over by the early enhancer EEα (Element A) in mammals. Another hypothesis is that element E originated as an alternative promoter. In this case, during the co-transcriptional splicing, element E would be in close proximity to the main promoter (looping) leading to increase local concentration of transcriptional cofactors and therefore enhanced transcription from the main promoter – the role and mechanism, as we understand it today, attributed to enhancers. As such, our data, may present a novel evolutionary mechanism for the origin of some enhancers.

Element A is uniquely present and strongly conserved in all mammals. It includes the EEα enhancer that has been shown experimentally to be required for early expression of *POU1F1* (10,12,13), although the conserved region of element A is much bigger than EEα. Extensive Blast searches of mammalian and non-mammalian genomic and other sequence databases did not reveal any sequence clearly related to element A other than upstream of the *POU1F1* gene in mammals. In early developing pituitary, element A is occupied by the giant platform-like protein ATBF1, that through its multiple DNA- and protein-binding domains consolidates multiple signals regulating the early expression of the gene (12), before POU1F1 transcription factor takes over its own regulation in autoregulatory fashion. ATBF1 is expressed only during a very short window of pituitary and brain development and only in a very limited number of tissues, raising the possibility that its role in the pituitary is to keep the *POU1F1* regulatory region available for regulation until appearance of POU1F1 to take over the transcriptional regulation of its locus.

The availability of genomic sequences from several hundred vertebrate species provides an abundance of data for comparative studies of the type presented here. The approach is inevitably broad-brush, partly because the data are sometimes incomplete and also because of intrinsic limitations. The main novel observation made in this study concerns the switch which occurred with the emergence of mammals in evolution. This transition involved the acquisition of additional elements in the *POU1F1* regulatory region, which potentially gave rise to a superenhancer, simultaneously losing the alternative upstream exon and promoter activity. The switch seems to involve a change from use of an alternative promoter/exon as a significant feature in regulation of *POU1F1* transcription to regulation based solely on a number of enhancers. Such a change seems to be substantial, but it may in part reflect the rather close relationship between promoters and enhancers which is now well recognized (1,5).

The functional implications of this regulatory switch remain ambiguous. The overall role of POU1F1 seems similar in most vertebrates, and transcription levels in the pituitary are high throughout. A biological function specific to mammals could involve the regulation of prolactin with its role in controlling lactation. In the female mammal, prolactin secretion undergoes large variations associated with pregnancy, lactation and post-lactation involution of the mammary gland. These changes may involve recruitment of additional lactotrophs in preparation for lactation, and subsequent reversal of the process - involving substantial cellular plasticity (27). Involvement of POU1F1 in these processes could require transcriptional controls additional to those needed for development of the pituitary gland and maintenance of high expression of growth hormone, prolactin and thyrotropin. This need may have been met by acquisition of additional enhancer elements (A and D) with subsequent modification involving loss of the alternative upstream exon and in some cases elements E and/or D. It is also possible that acquisition of the additional strong regulatory element (A) in the region open early during the development of the pituitary was necessary to prevent heterochromatinization of the locus in brain development in mammals.

In sum, this study has followed the evolution of the regulatory region of the *POU1F1* gene and found a potential sequence of events that led to the self regulatory, superenhancer-like region maintaining high and reliable expression of the gene in the adult mammalian pituitary. To our knowledge, this is the first analysis of the evolution of enhancer-rich regulatory region in vertebrates. Our data not only suggest how an alternative promoter might become an enhancer but also underline the potential importance of maintenance of old regulatory elements in parallel with introduction of new elements in order to maintain gene expression and function.

## Materials and Methods

### Sources of pituitaries

Tissues from guinea fowl (*Numida meleagris*), alligator (*Alligator mississipiensis*) and bullfrog (*Lithobates catesbeianus*) were a kind gift from Drs Daley and Azizi laboratories at UCI. Mouse colony (C57Bl6) is established in D. S-K laboratory. All procedures are performed under approved protocols.

### Genomic sequences

Genomic sequences for *POU1F1* were extracted from the ensembl or NCBI wgs databases, following BLAST searching (28) to locate the genes. In some cases additional searches of NCBI databases were used to extend/complete genomic sequences. Details for species included in this paper are given in *SI Appendix,* **Table S1**.

### Transcriptomes

Pituitary transcriptomes for various mammals, birds, amphibia and fish were accessed through the NCBI SRA database, and used to assess transcription levels of *POU1F1*, and details of splicing patterns. Expression profiles were established as described previously (29).

### ATAC-seq

was performed as previously published (21). In brief, fresh tissue was re-suspended immediately in 1 ml ice cold nuclei permeabilization buffer (5%BSA, 0.2% (m/v) NP40, 1mM DTT in PBS solution) with 1X complete EDTA-free protease inhibitor and homogenized using a syringe and needle followed by slow rotation for 10 minutes at 4°C. The nuclei suspension was then filtered through a 40µM cell strainer and centrifuged for 5 minutes at 500xg at 4℃. The nuclei pellet was resuspended in an ice cold 50 µl tagmentation buffer. Nuclei concentration was adjusted to 2,000-5,000 nuclei/µl and 10 µl of the suspension was used for tagmentation. 0.5 µl Tagment DNA Enzyme 1 (FC-121-1030, Illumina) was added to the 10µl suspension. The reaction mix was thoroughly pipetted and incubated 30 minutes with 500 rpm at 37 °C. After the tagmentation reaction completed, the DNA was isolated using Qiagen PCR Purification Kit (Cat.#.28304, Qiagen) and eluted in 20 µl Elution Buffer. The eluted DNA fragments were then amplified by PCR with Nextera compatible indexed sequencing i5 and i7 adapters using NEBNext 2x PCR Master Mix PCR kit (M0541, NEB). The amplified DNA library was fragment size selected from 200bp to 800bp using Ampure XP beads (A63880, Beckman Coulter). The quality of the ATAC-Seq libraries was assessed by Agilent 2100 bioanalyzer (Agilent Technologies, Inc.). ATAC-Seq libraries was pooled and run on a NovaSeq 6000 System (Flow Cell Type S4) Illumina sequencer with a paired-end read of 100 bp to harvest about 50 million paired-end reads per sample.

### Identification of enhancer elements and POU1F1 binding sites

In many cases conserved enhancer elements were identified upstream of the *POU1F1* gene in various species by BLAST searching of wgs databases with sequences from related organisms (see example in *SI Appendix,* **Fig. S6**). In other cases more-focussed searches of sequences upstream of the *POU1F1* gene were necessary, using BLAST or clustalw (30). Potential POU1F1 binding sites were identified using PROMO (31), with taxon restriction to Craniata.

### Sequence alignments

Alignment of enhancer sequences was carried out using M-Coffee (32,33) followed by manual adjustment and evaluation using the TCS procedure (34).

## DATA AVAILABILITY

Data is deposited into the GEO database and the GSE number will be shared upon publication of the manuscript.

## ACKNOWLEDGMENTS

Authors would like to thank Drs. Monica Daley and Manny Azizi at UCI and members of their laboratories in obtaining pituitaries from different species. Work in Dorota Skowronska-Krawczyk laboratory is partially supported by unrestricted RPB grant to the University of California, Irvine, Department of Ophthalmology.

## CONTRIBUTION

D. S-K and M.W. - Conceptualisation, data analysis, writing, editing, funding; Q.X. – experiments, data analysis, editing; M.K. – data analysis, editing;

---

**Supplementary Table S1.**
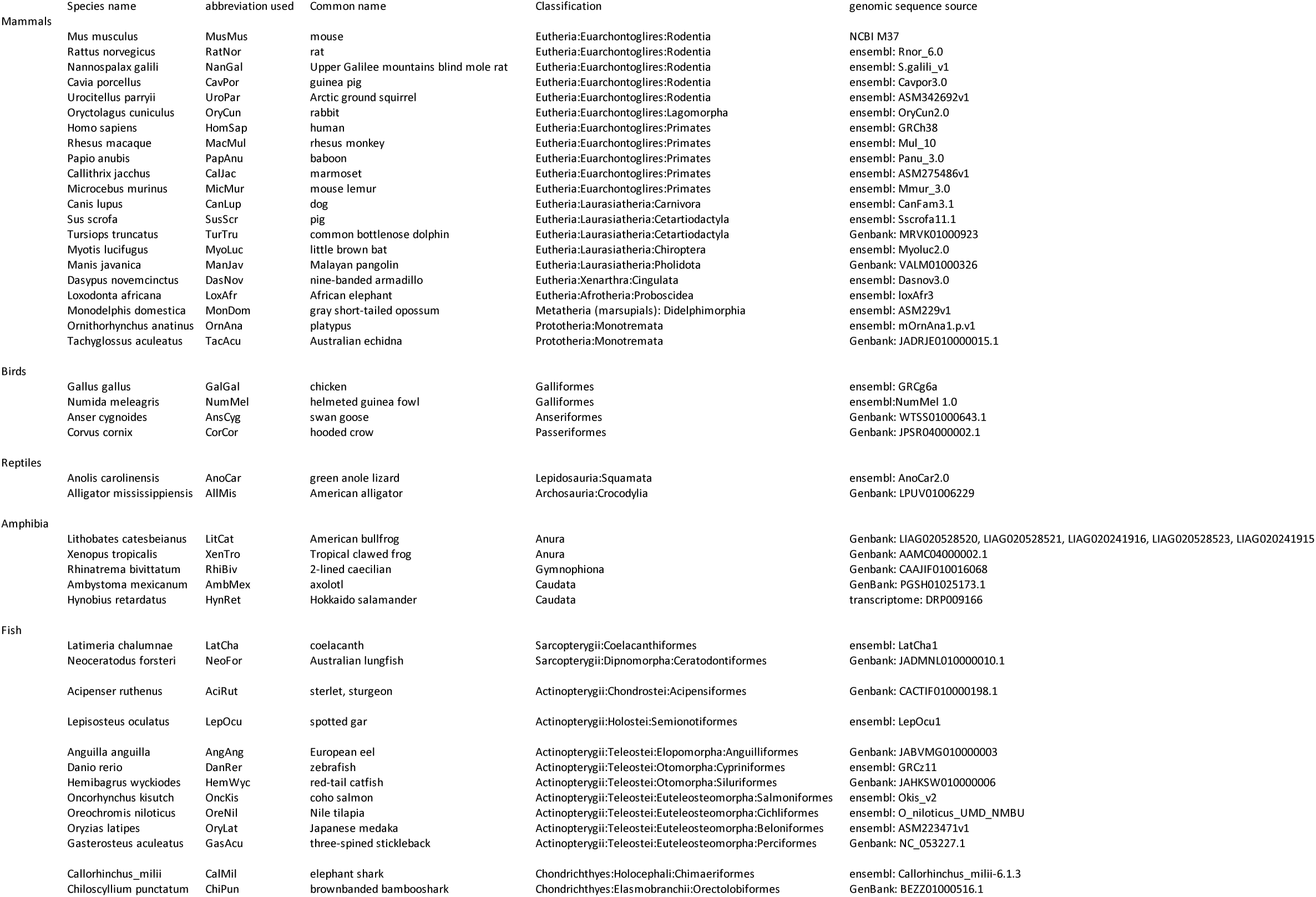
Species studied and sources.

**Supplementary Table S2.**
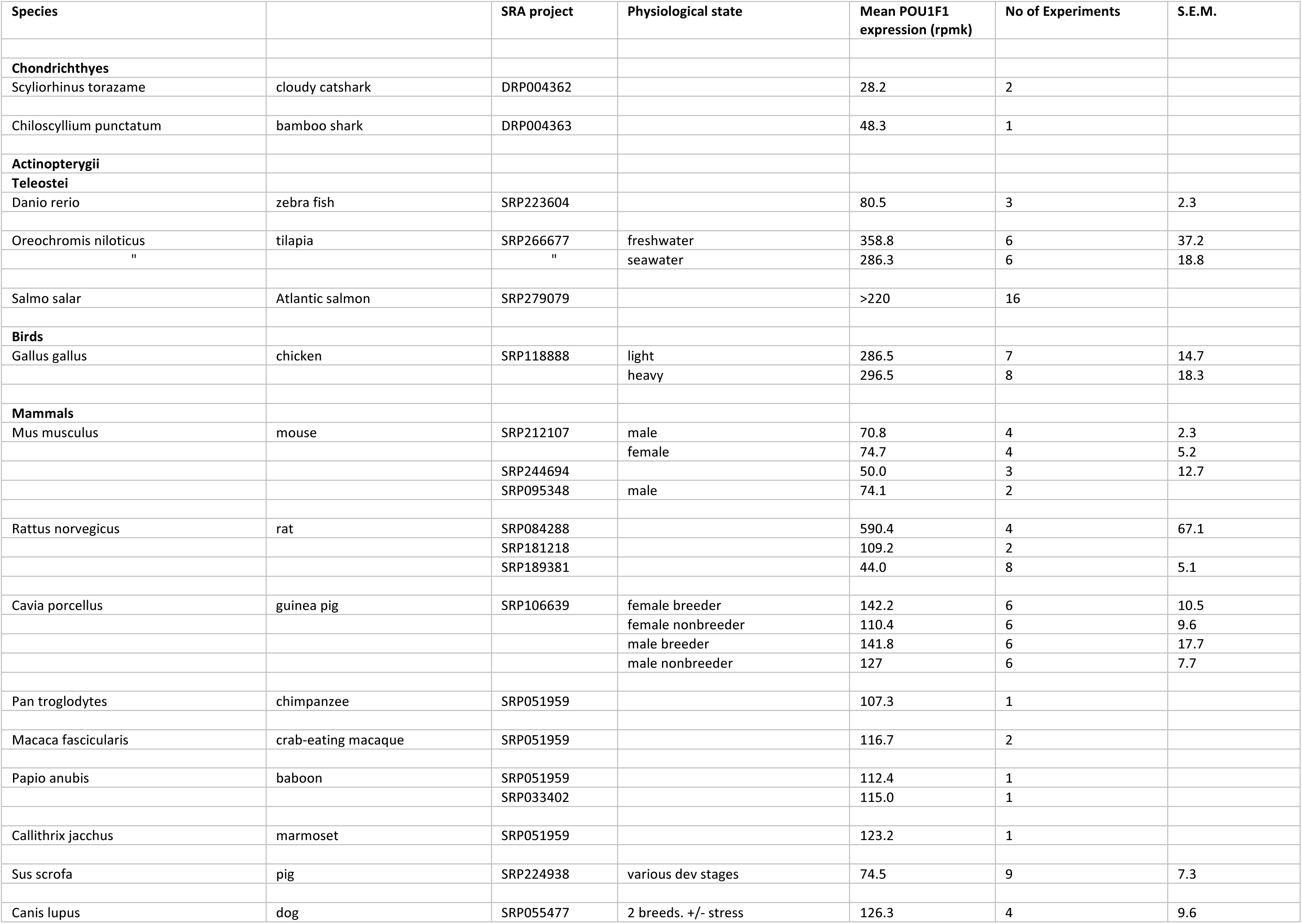
Expression of POU1F1 in pituitary transcriptomes from various species.

**Supplementary Fig. S1.**
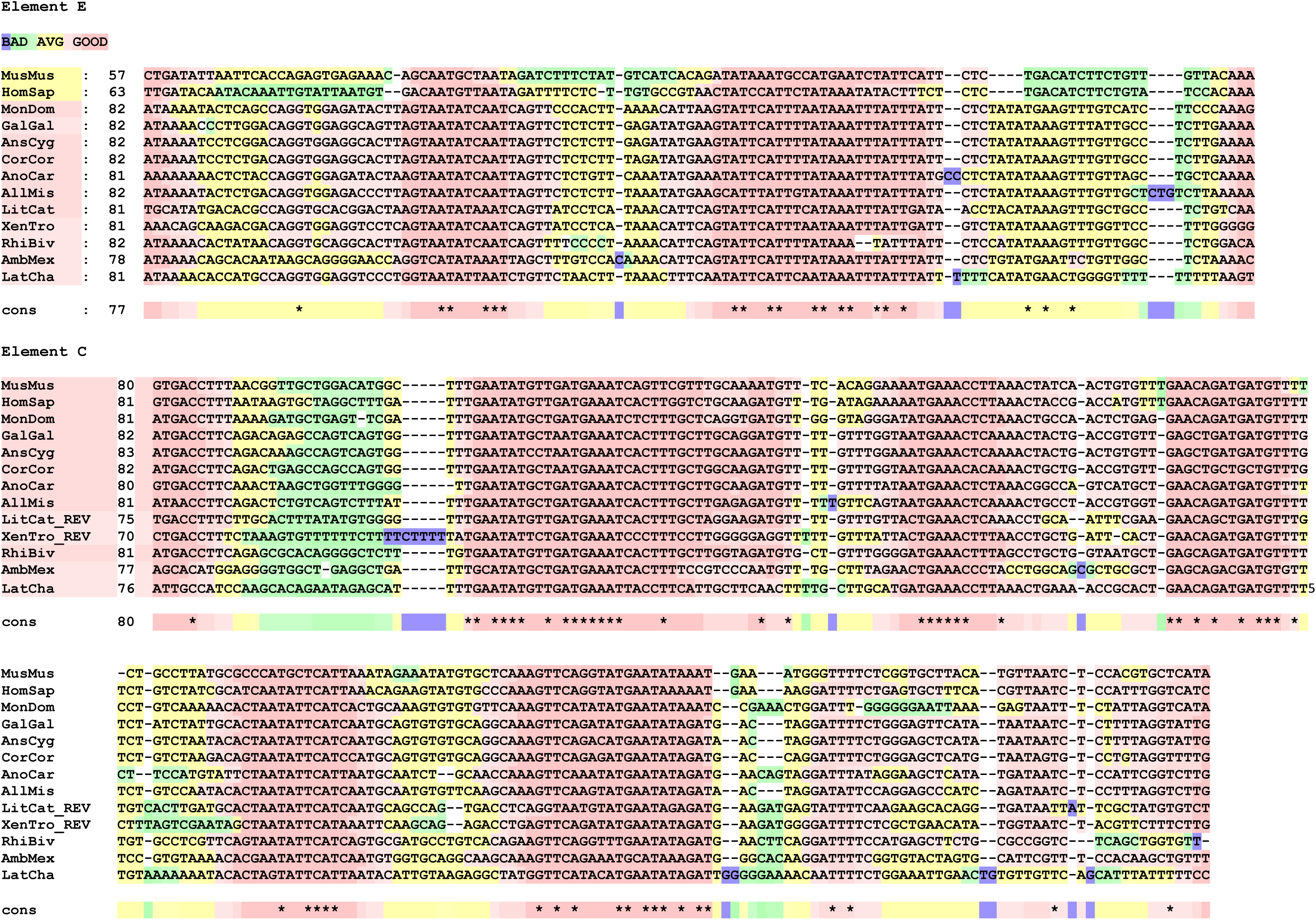
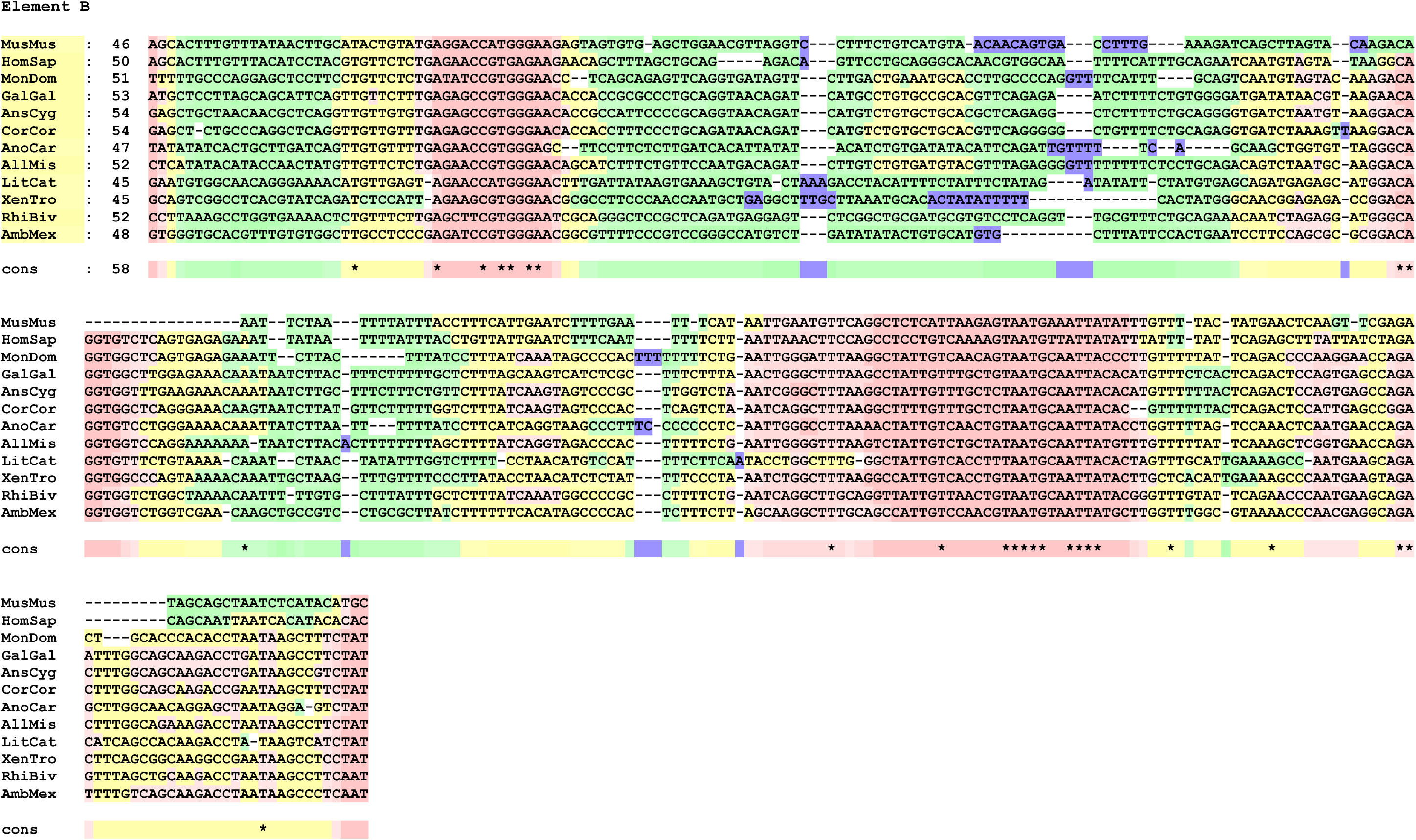
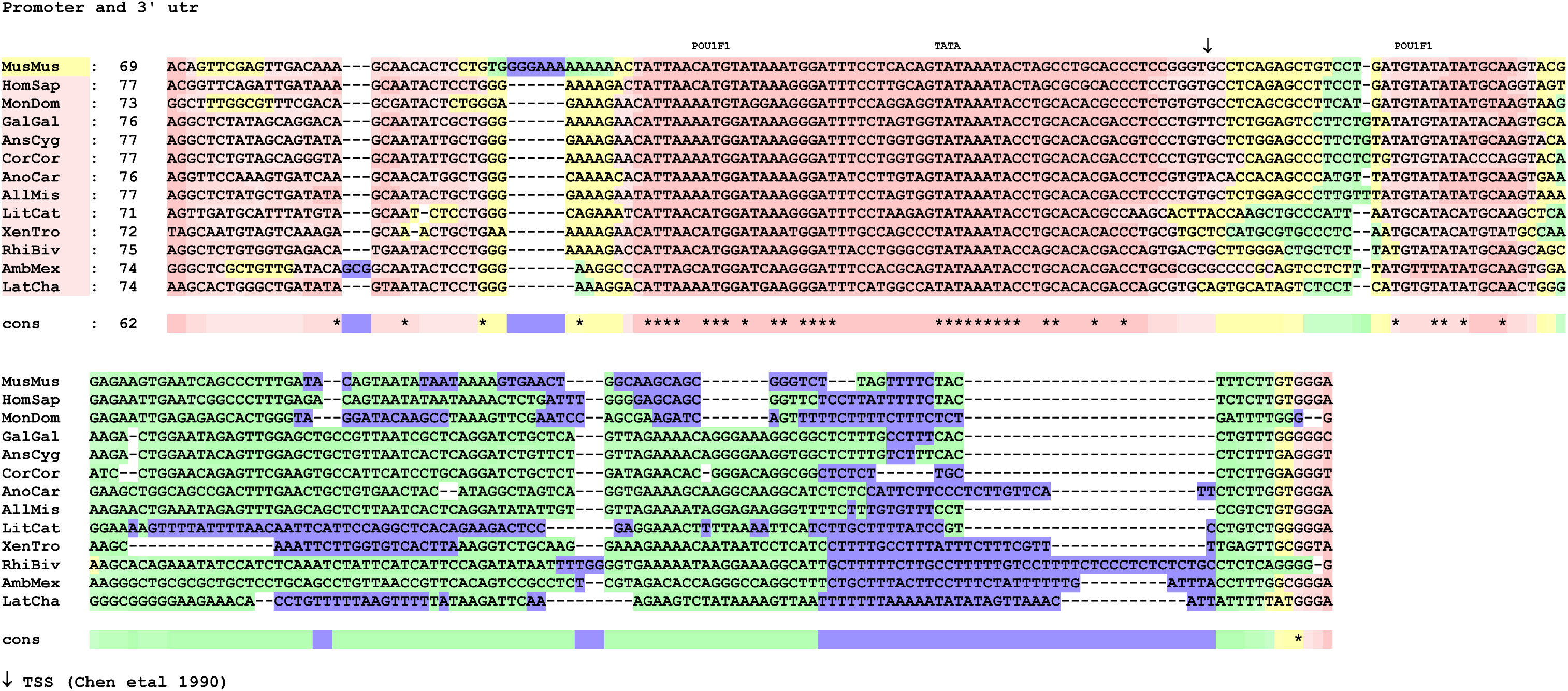
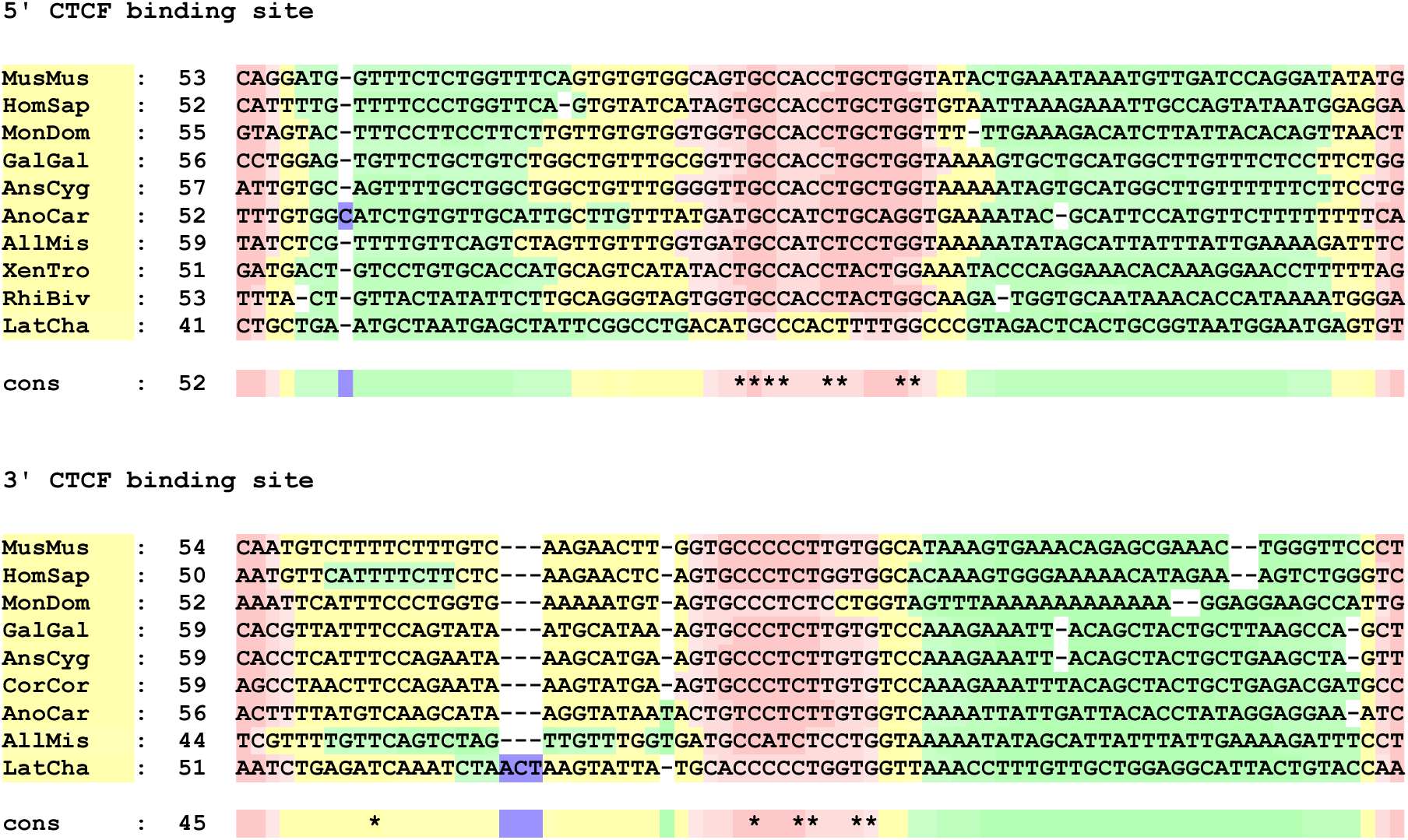
Sequence alignments for POU1F1 enhancer elements E, C and B and promoter/utr from a range of tetrapods. Alignments were performed using the M-Coffee procedure (32,33) followed by manual adjustment and evaluation using the TCS procedure (34). ↓ indicates the transcription start site, based on Chen et al (1990) (8). Alignments of 5’ and 3’ CTCF binding sites are also included. Abbreviations of sequence names are: MusMus, mouse; HomSap, human; MonDom, oppossum (mammals); GalGal, chicken; AnsCyg, swan goose; CorCor, hooded crow (birds); AnoCar, anole lizard; AllMis, American alligator (reptiles); LitCat, bullfrog; XenTro, Western clawed frog; RhiBiv, 2-lined caecilian; AmbMex, axolotl (amphibia); LatCha, coelacanth (sarcopterygian fish, included for comparison).

**Supplementary Fig. S2.**
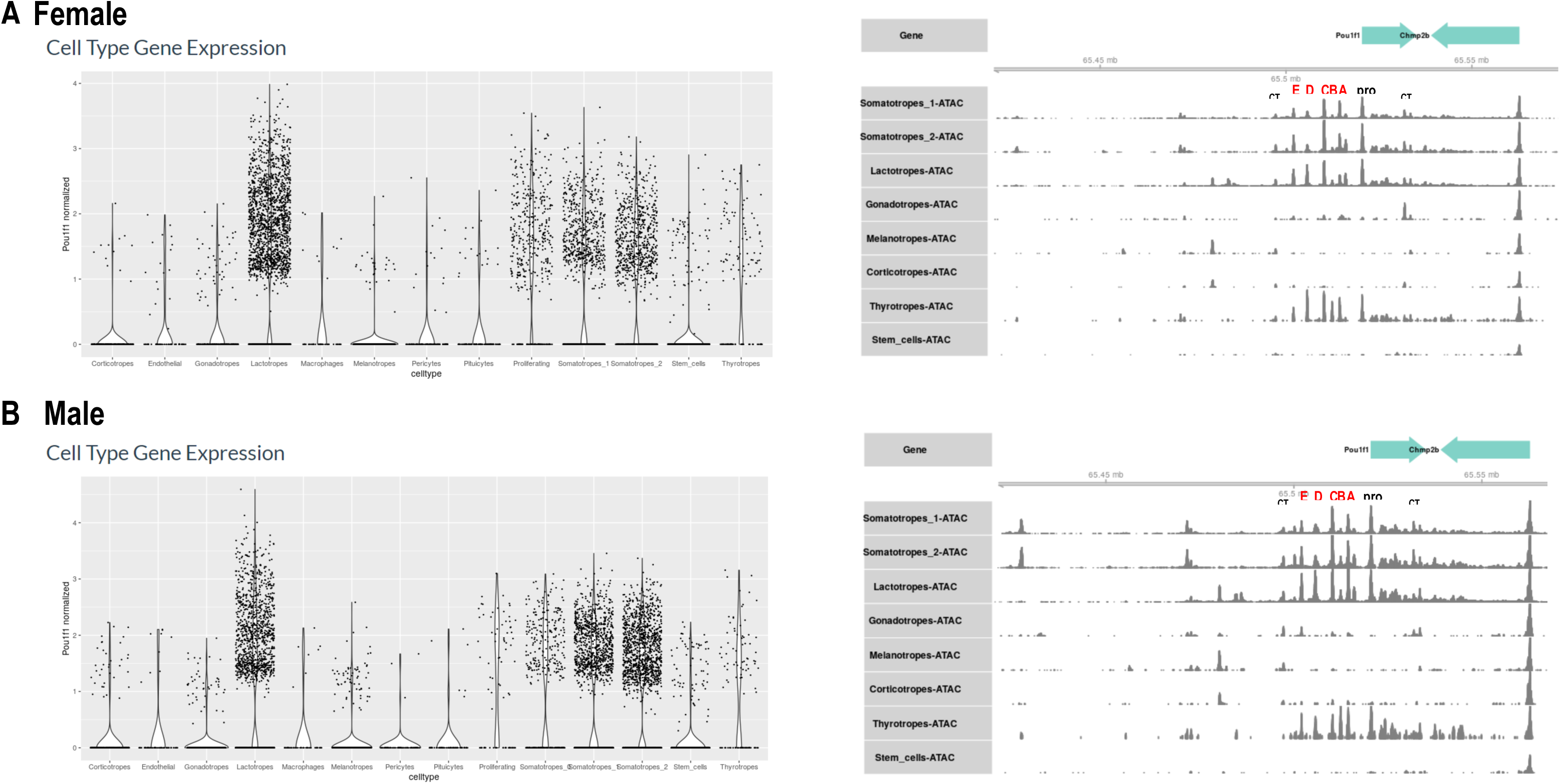
Single cell pituitary multi-omics analysis (15) reveals the DNA accessibility of conserved regulatory elements in every POU1F1 positive cell type.

**Supplementary Fig. S3.**
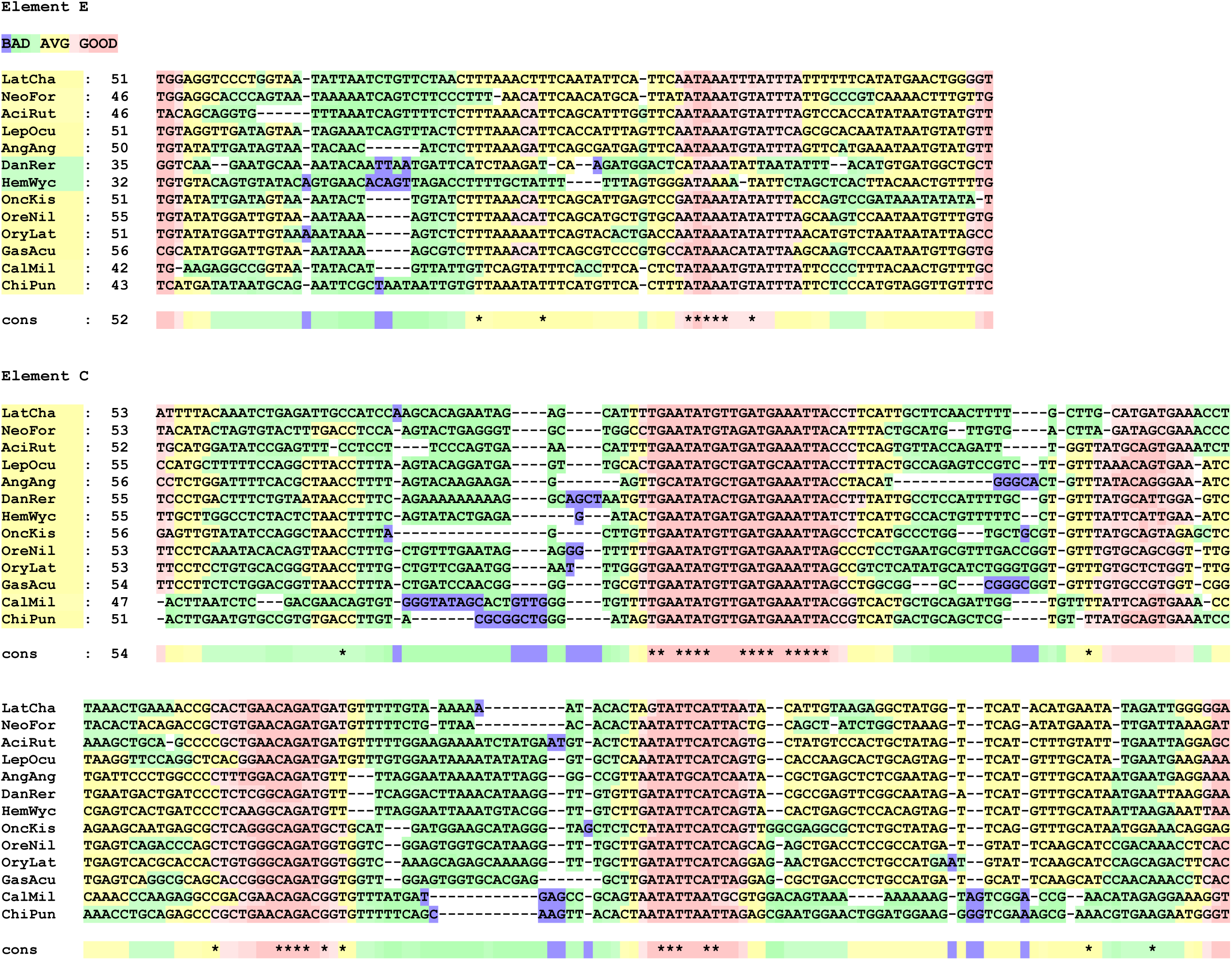
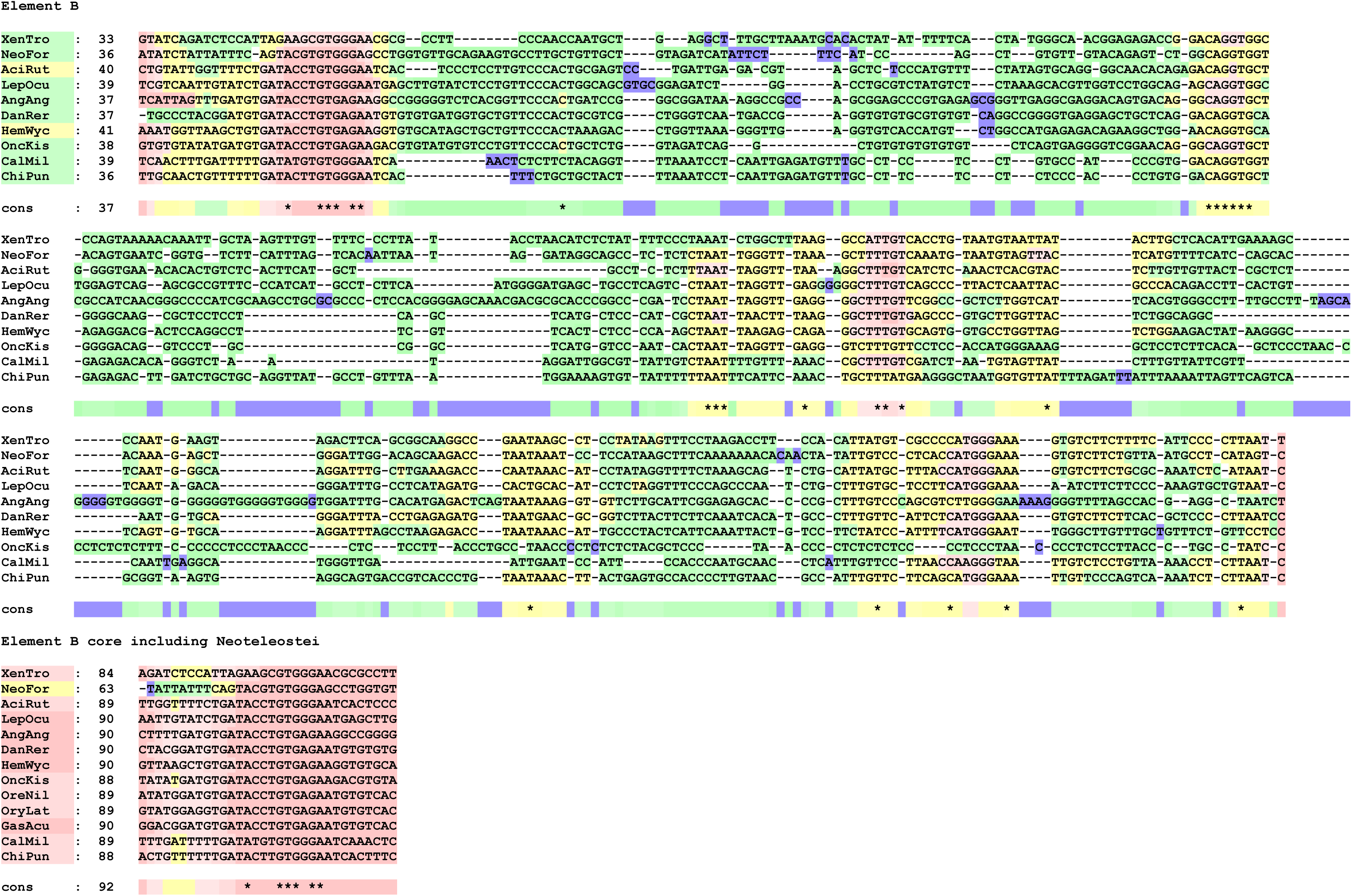
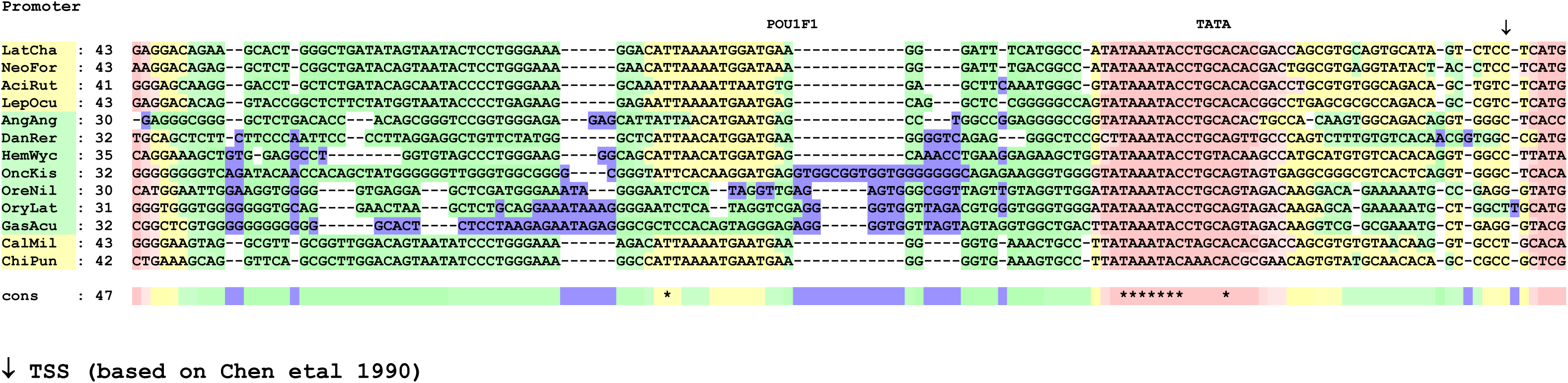
Sequence alignments for POU1F1 enhancer elements E, C and B and promoter from a range of fish. Alignments were performed using the M-Coffee procedure (32,33) followed by manual adjustment and evaluation using the TCS procedure (34). ↓ indicates the transcription start site, based on Chen et al (1990) (8). Abbreviations of sequence names are: LatCha, coelacanth; NeoFor, Neoceratodus forsteri, Australian lungfish (sarcopterygian fish); AciRut, sturgeon (Chondrostei); LepOcu, spotted gar (Holostei); AngAng, European eel (Elopomorpha, Teleostei), DanRer, zebra fish; HemWyc, red-tail catfish (Otomorpha, Teleostei), OncKis, coho salmon; OreNil, Nile tilapia; OryLat, Japanese ricefish; GasAcu, stickleback (Euteleosteomorpha, Teleostei); CalMil, elephant shark; ChiPun, bamboo shark (Chondrychthyes). See Supplementary Table 1 for full scientific names. For the Element B the coelacanth (LatCha) sequence is not available; the Xenopus (XenTro) sequence is included to provide a link to Supplementary Fig. 1. Euteleost sequences (OreNil, OryLat, GasAcu) are omitted from the full Element B alignment because they show similarity over only a short region; alignment of this region is shown as "Element B core".

**Supplementary Fig. S4.**
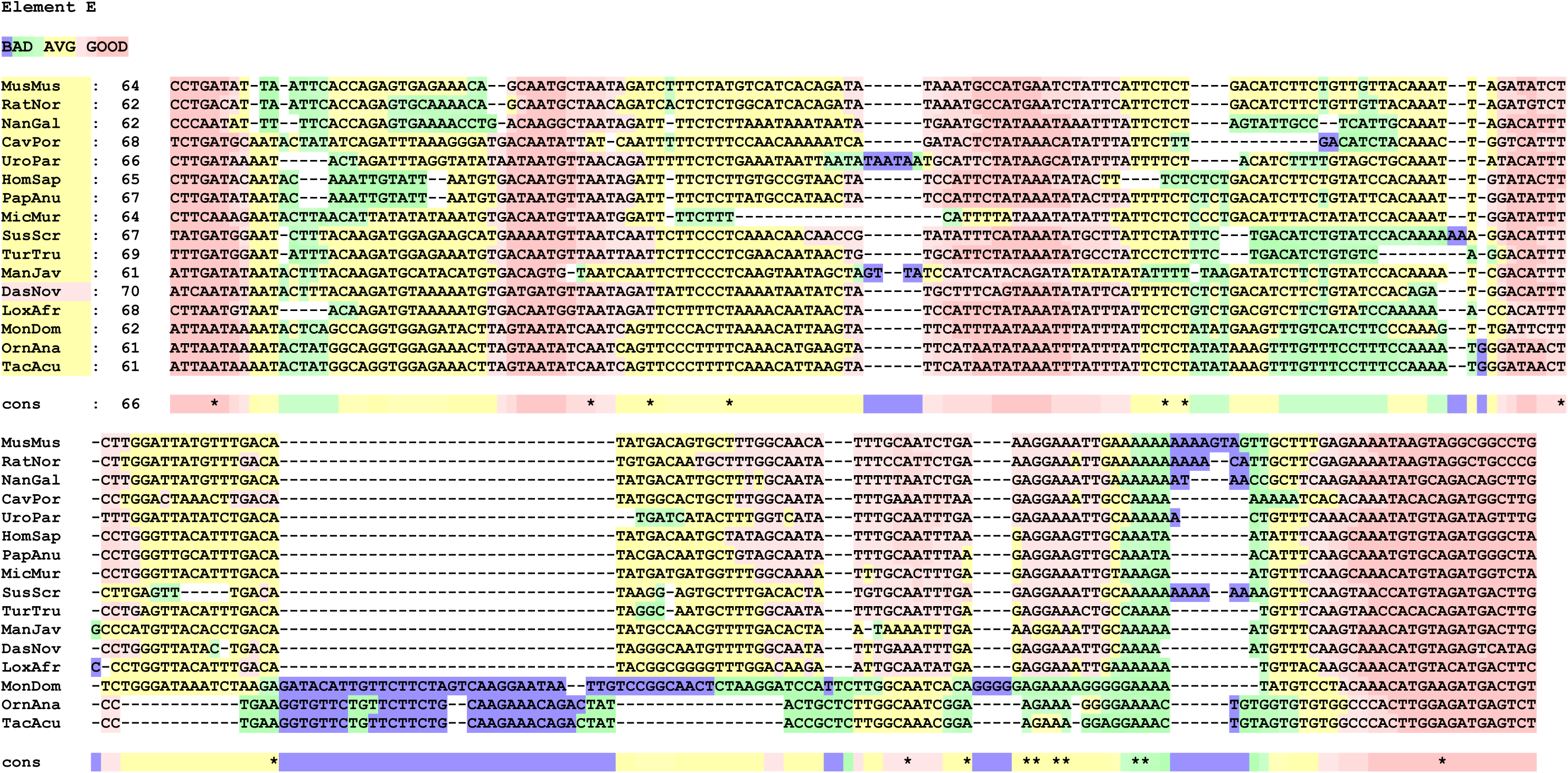
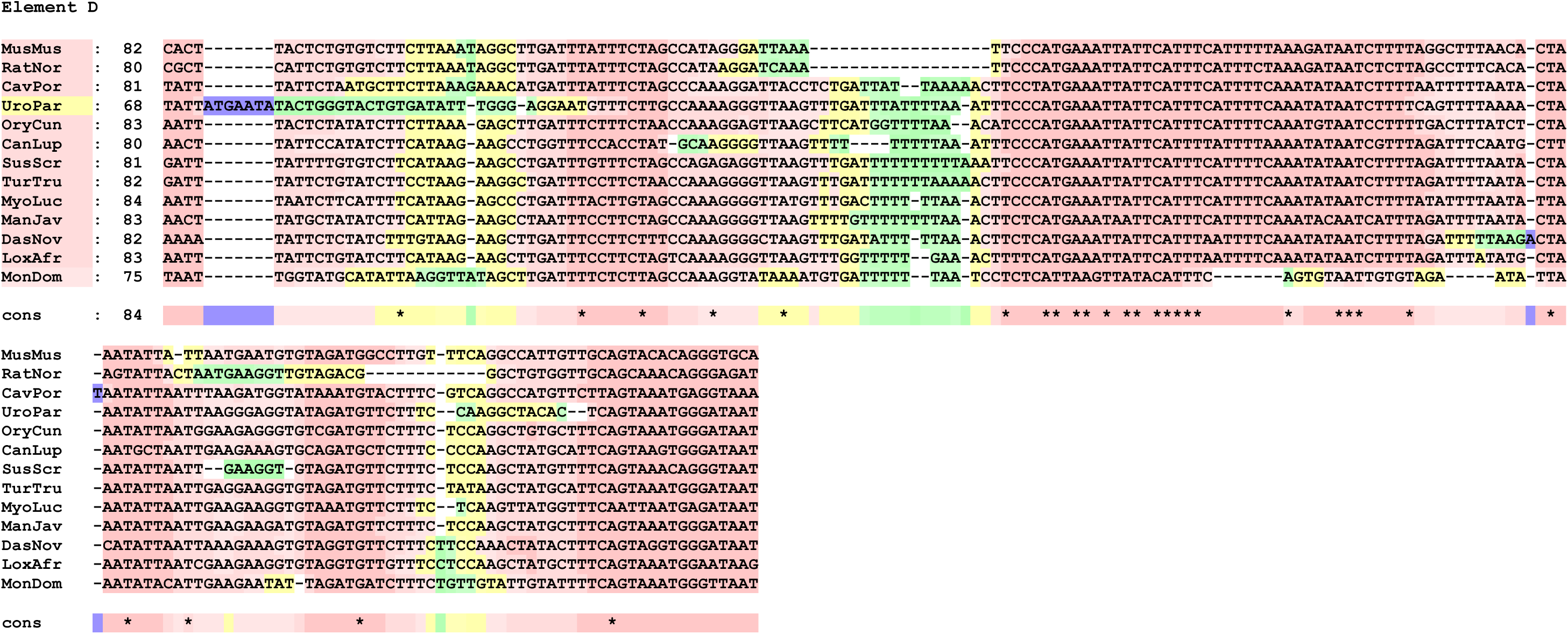
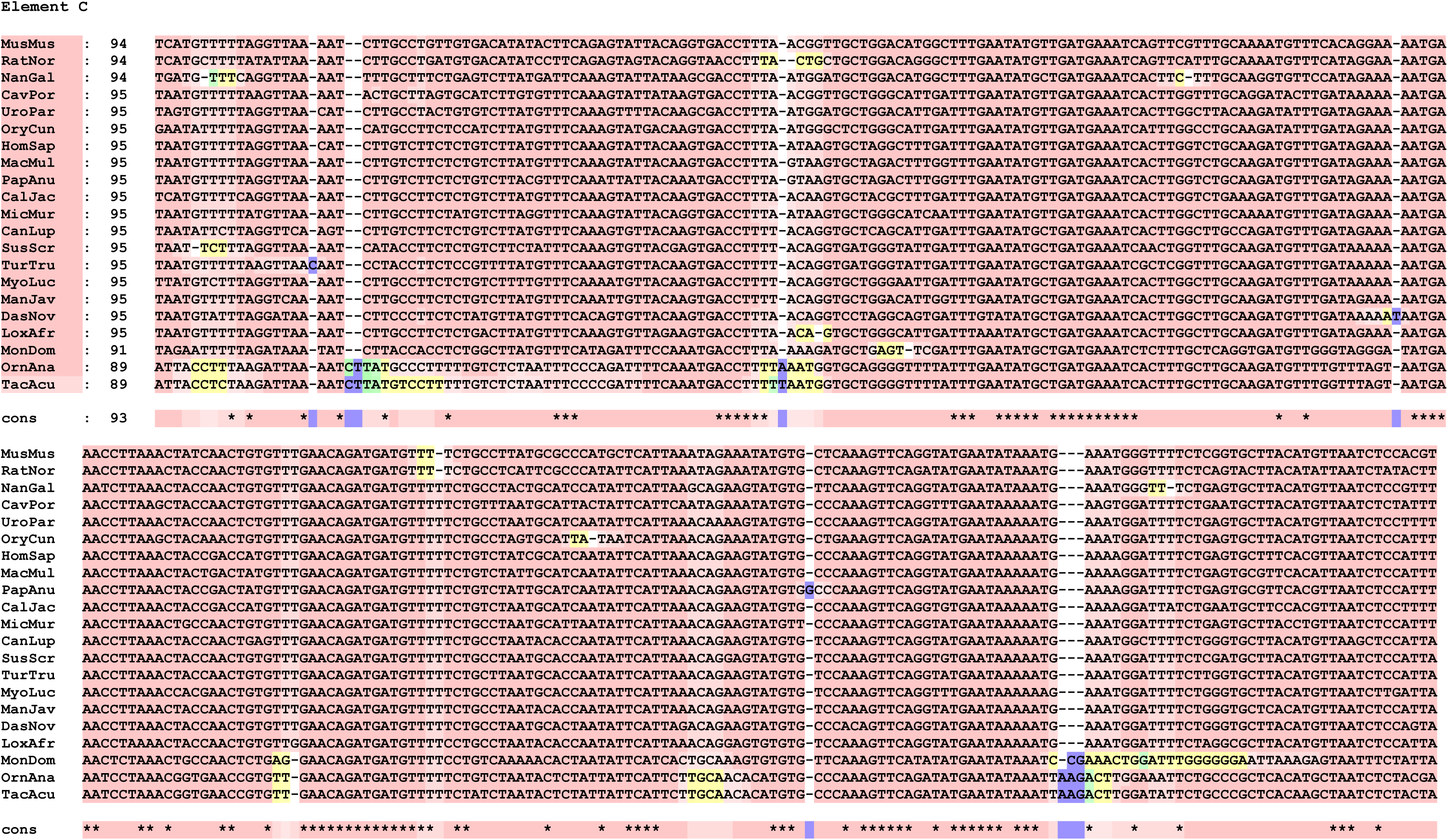
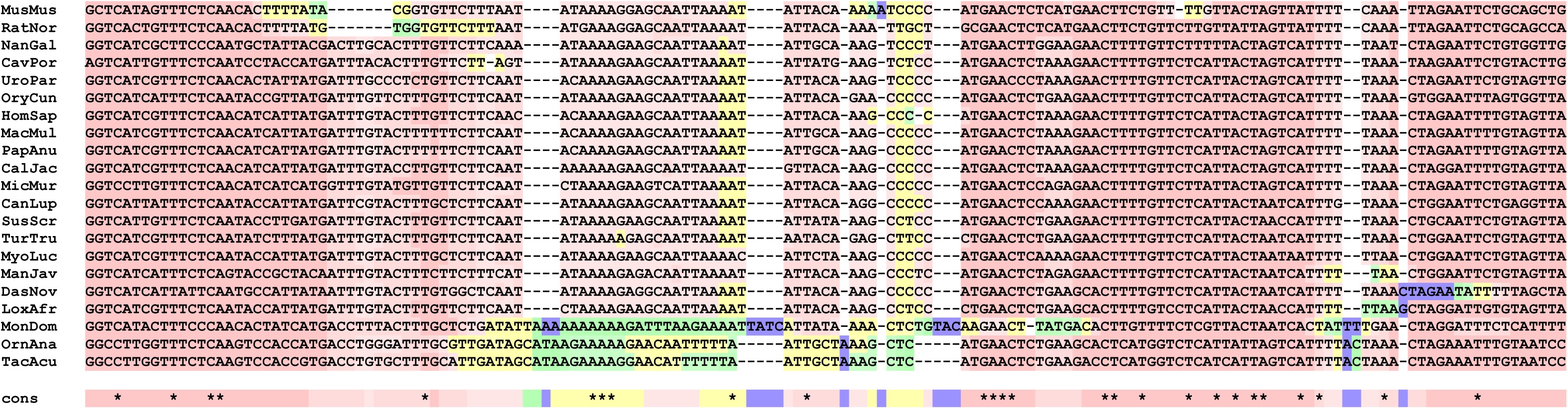
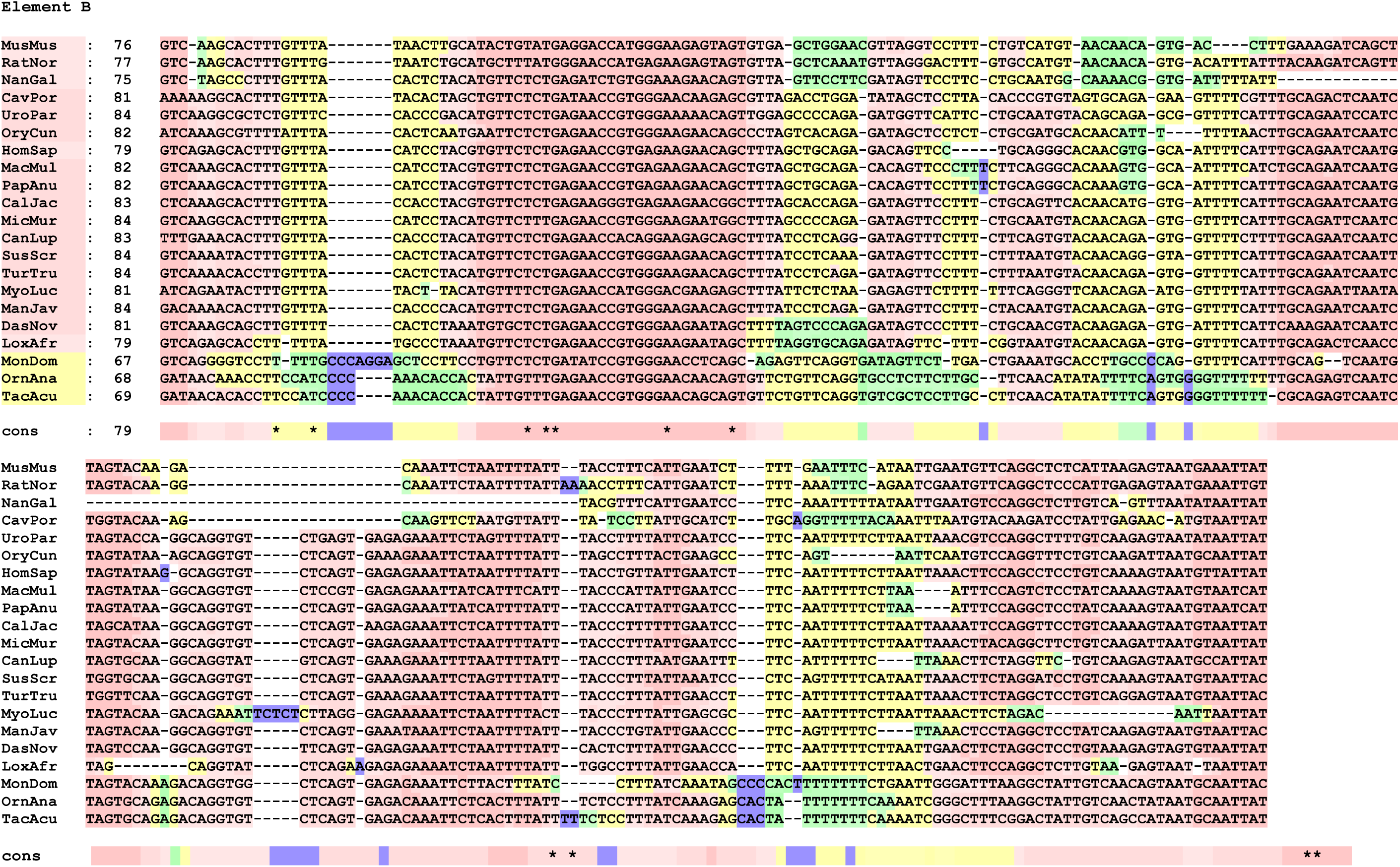
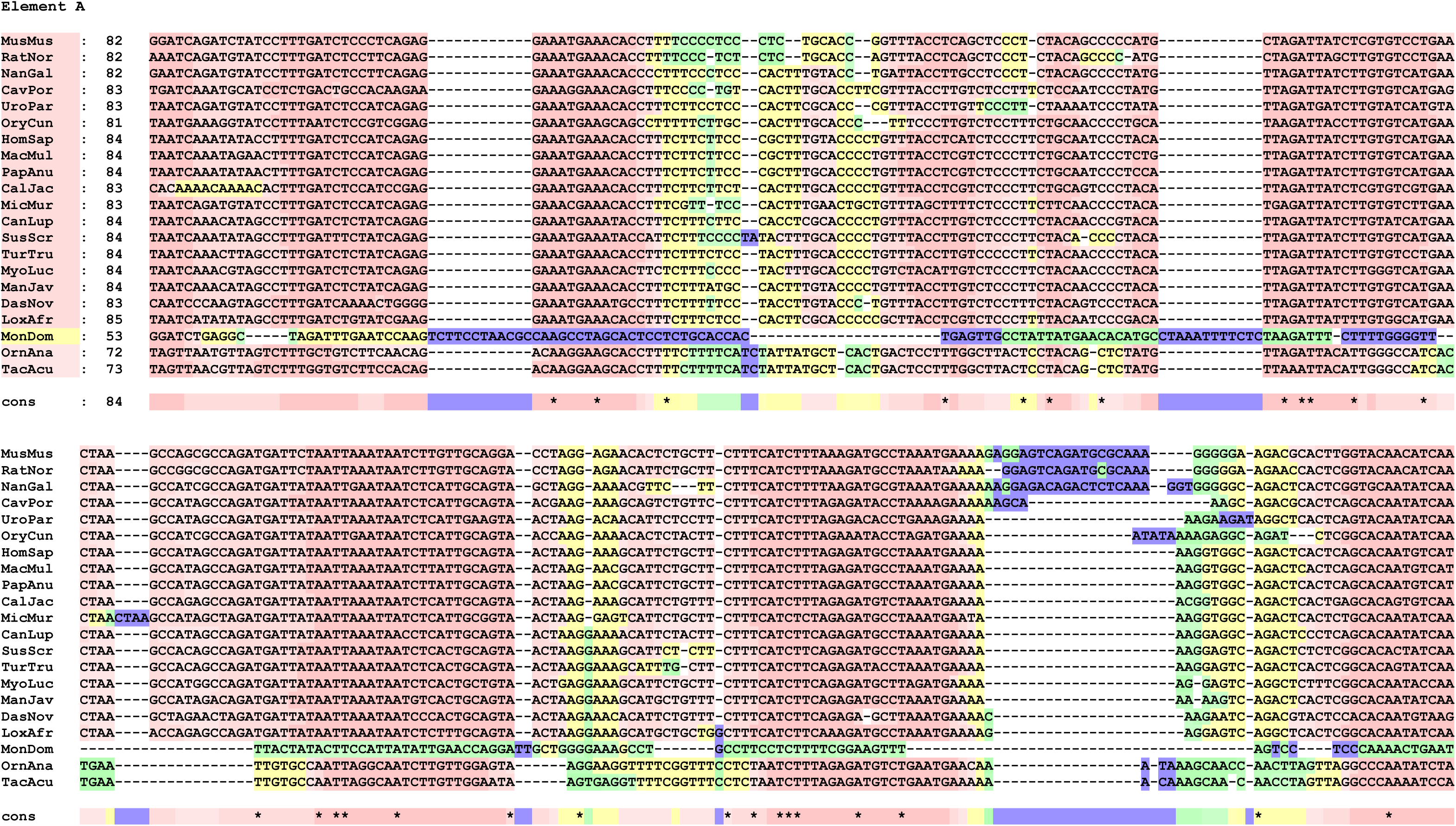
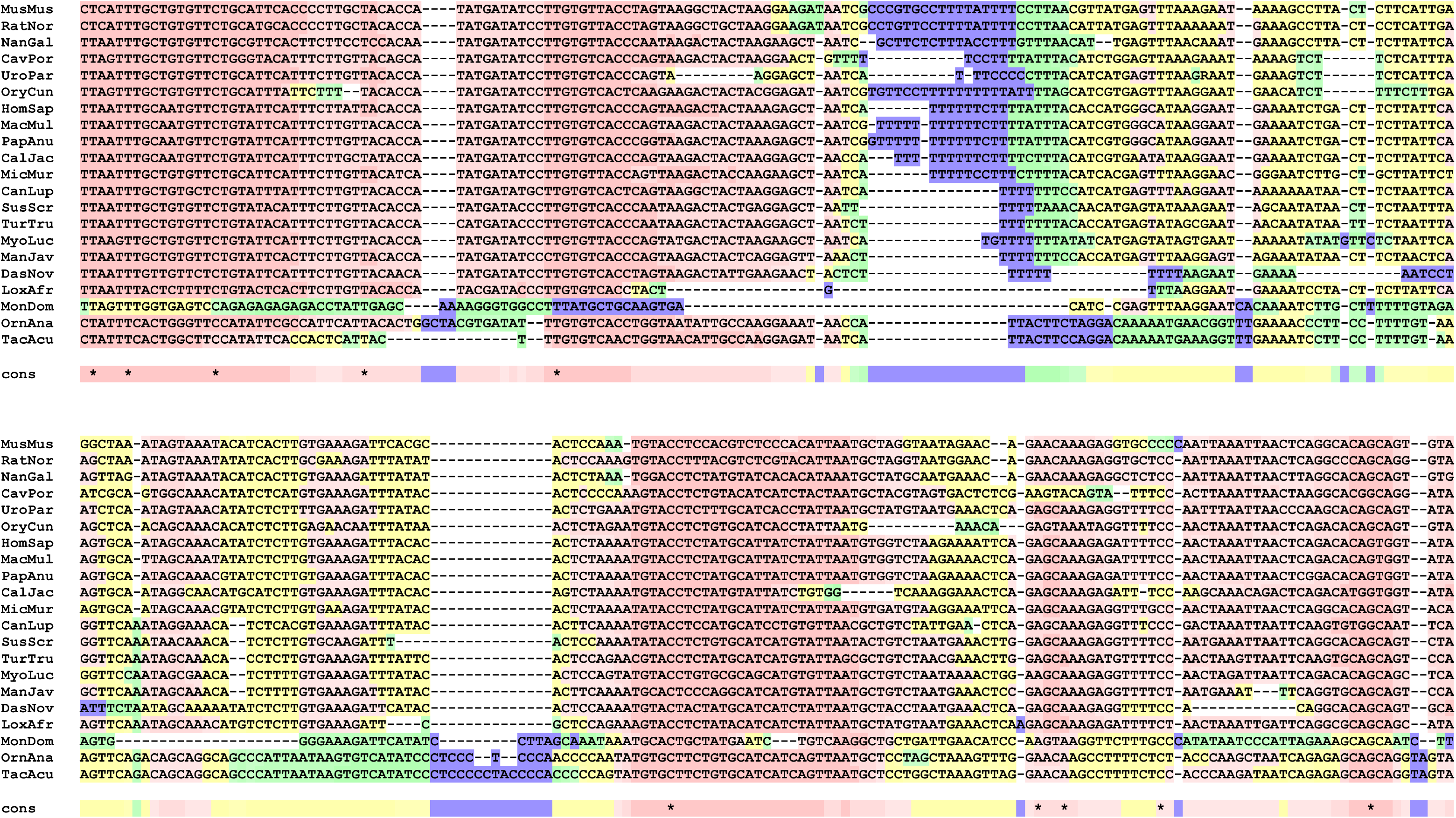
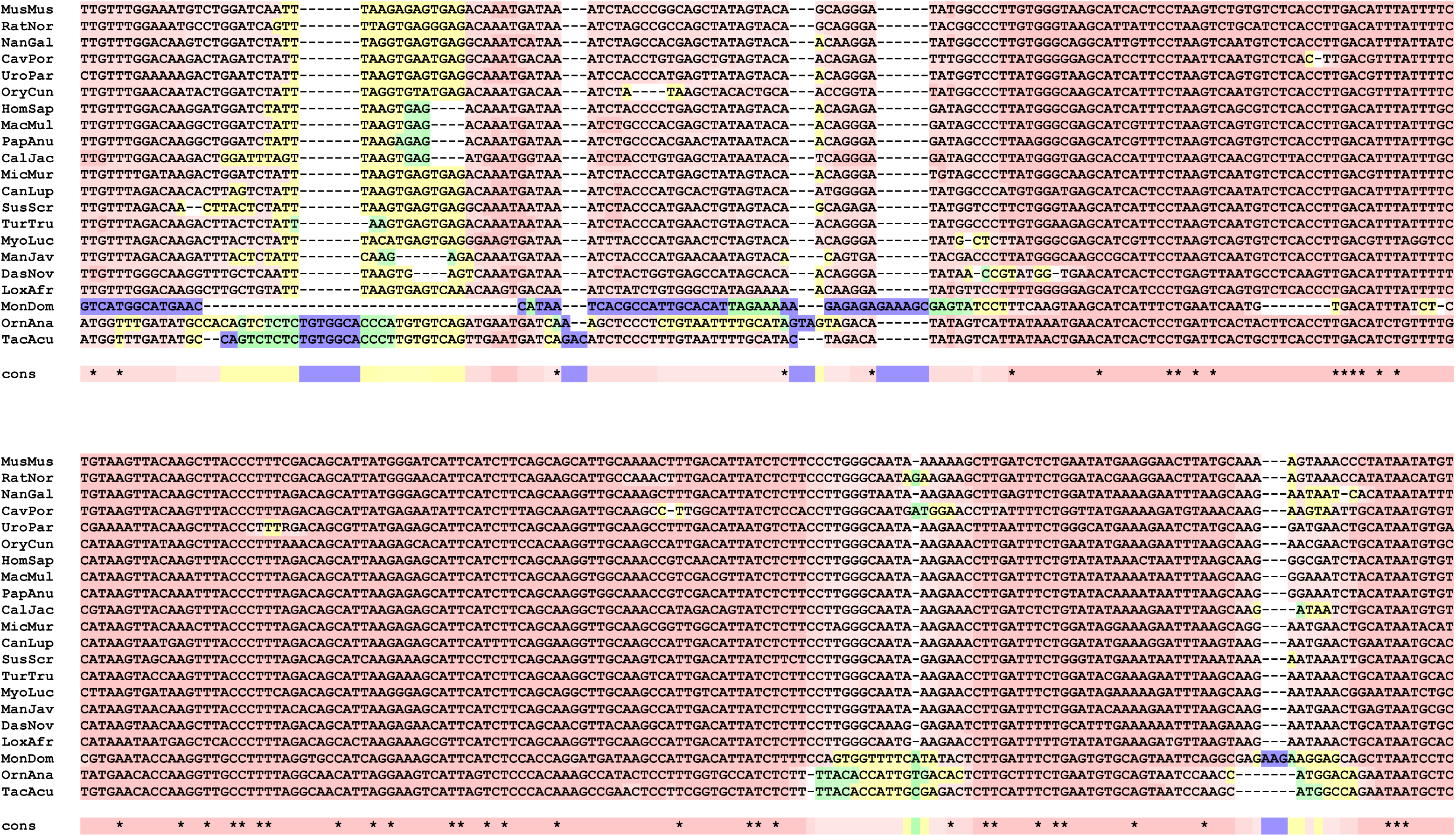
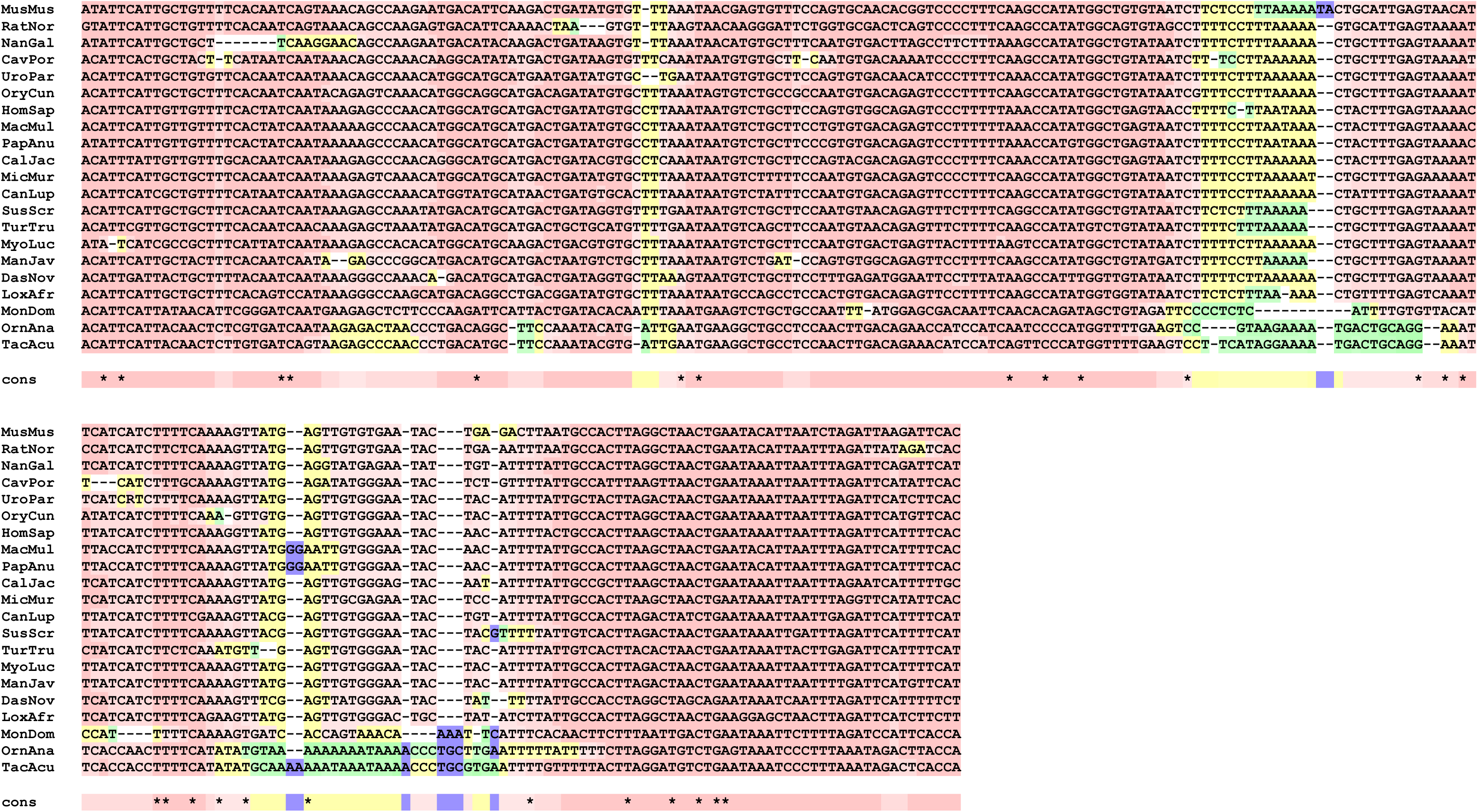
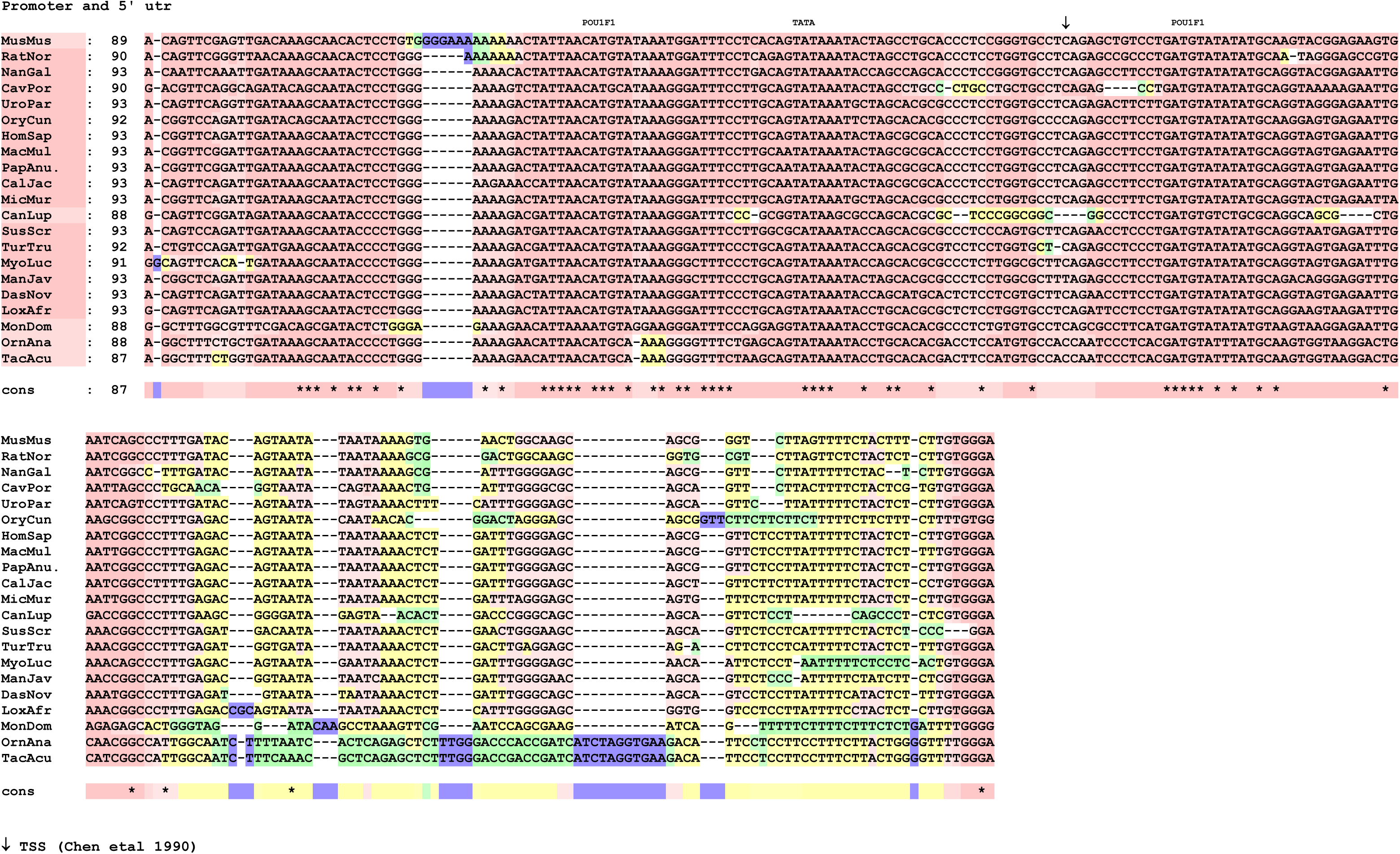
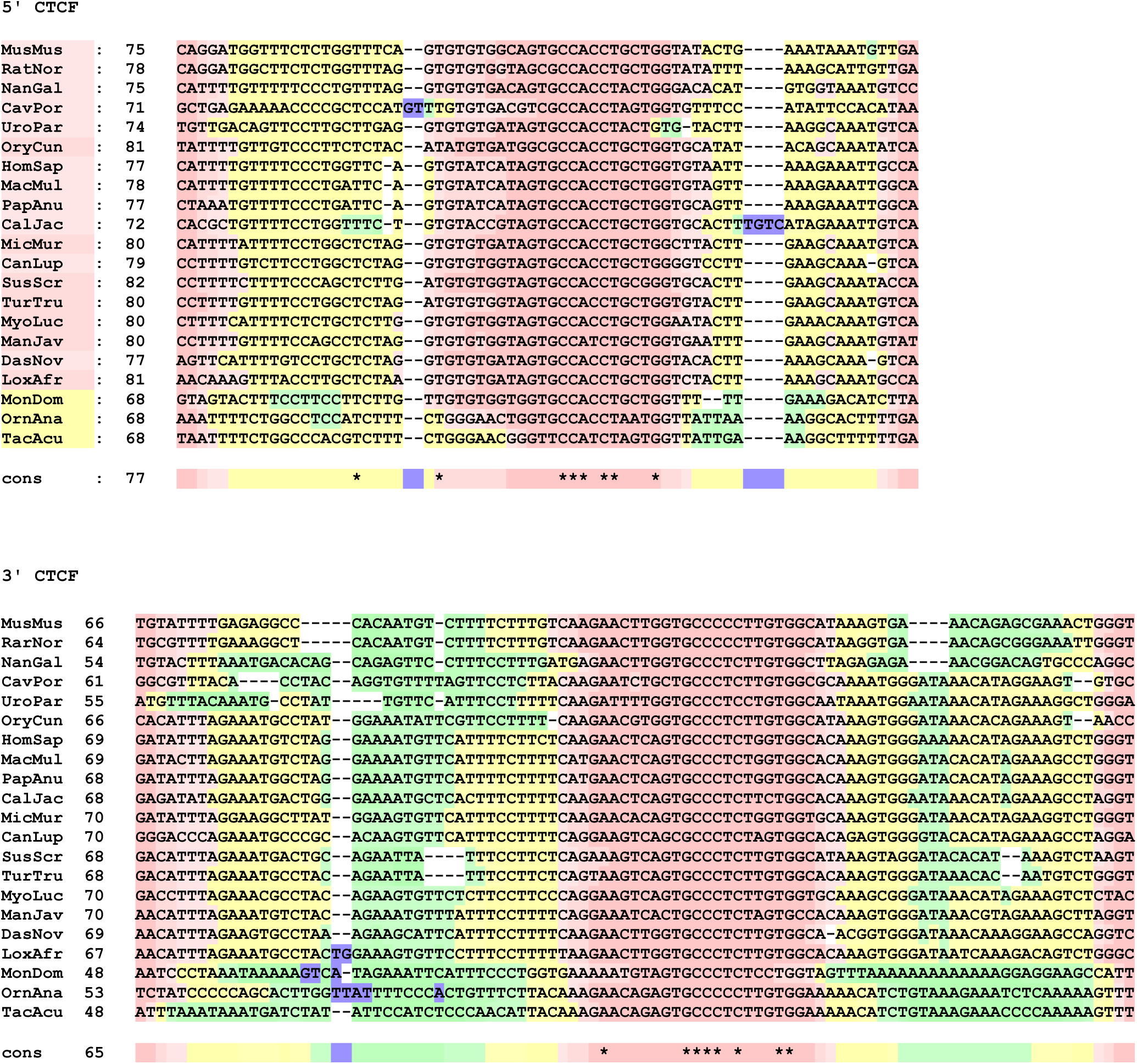
Sequence alignments for POU1F1 enhancer elements E-A and promoter/utr from a range of mammals. Alignments were performed using the M-Coffee procedure (32,33) followed by manual adjustment and evaluation using the TCS procedure (34). ↓ indicates the transcription start site, based on Chen et al (1990) (8). Abbreviations of sequence names are: MusMus, mouse; RatNor, rat; NanGal, Upper Galilee mountains blind mole rat; CavPor, guinea pig; UroPar, Arctic ground squirrel; OryCun, rabbit; HomSap, human; MacMul, rhesus monkey; PapAnu, baboon; CalJac, marmoset; MicMur, mouse lemur; CanLup, dog; SusScr, pig; TurTru, bottlenose dolphin; MyoLuc, little brown bat; ManJav, pangolin; DasNov, nine-banded armadillo; LoxAfr, African elephant; MonDom, oppossum; OrnAna, duck-billed platypus; TacAcu, echidna.

**Supplementary Fig. S5.**
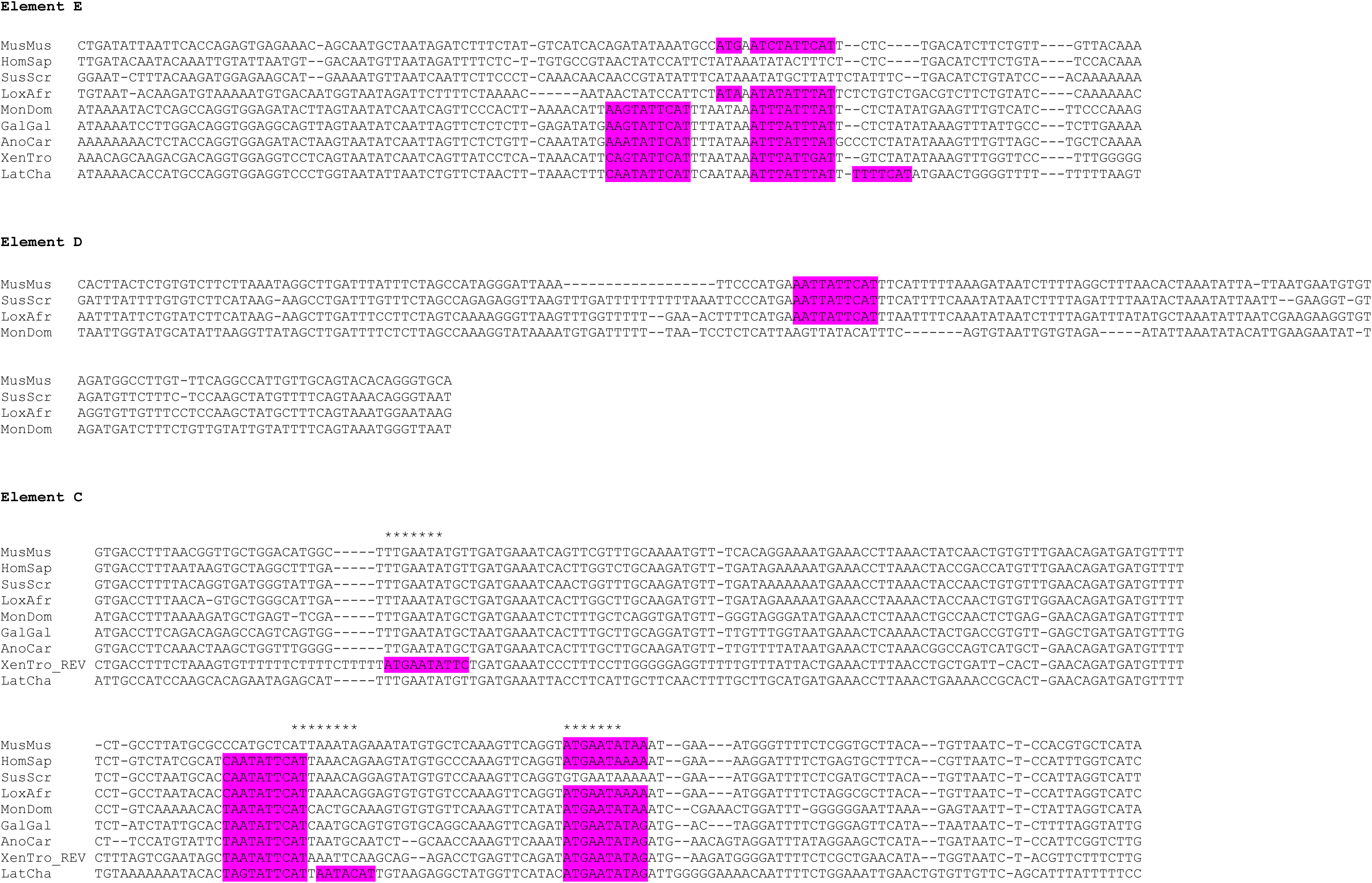
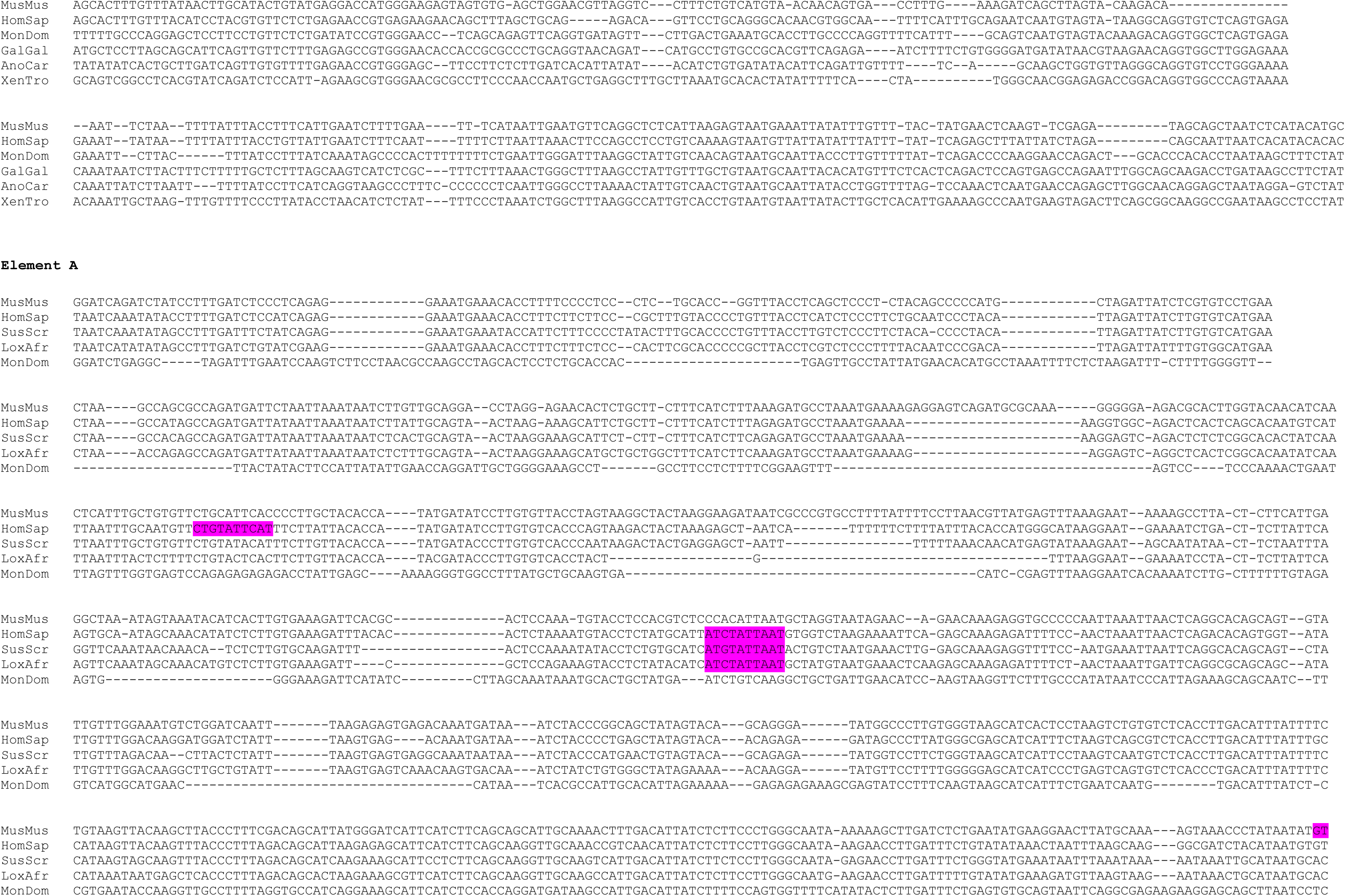
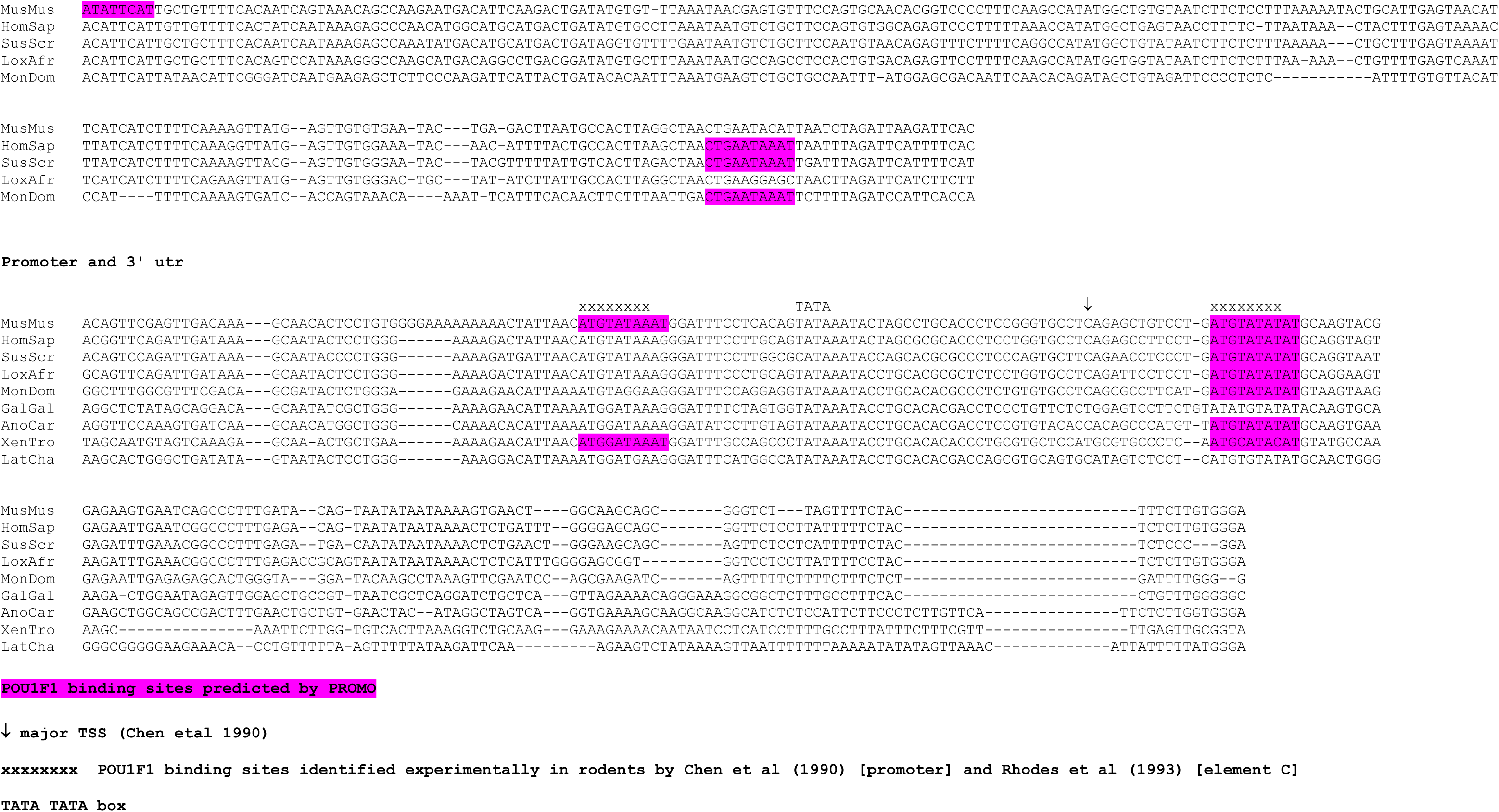
Alignments of selected Element E-A and promoter sequences, with positions of POU1F1 binding sites predicted by PROMO highlighted.

**Supplementary Fig. S6.**
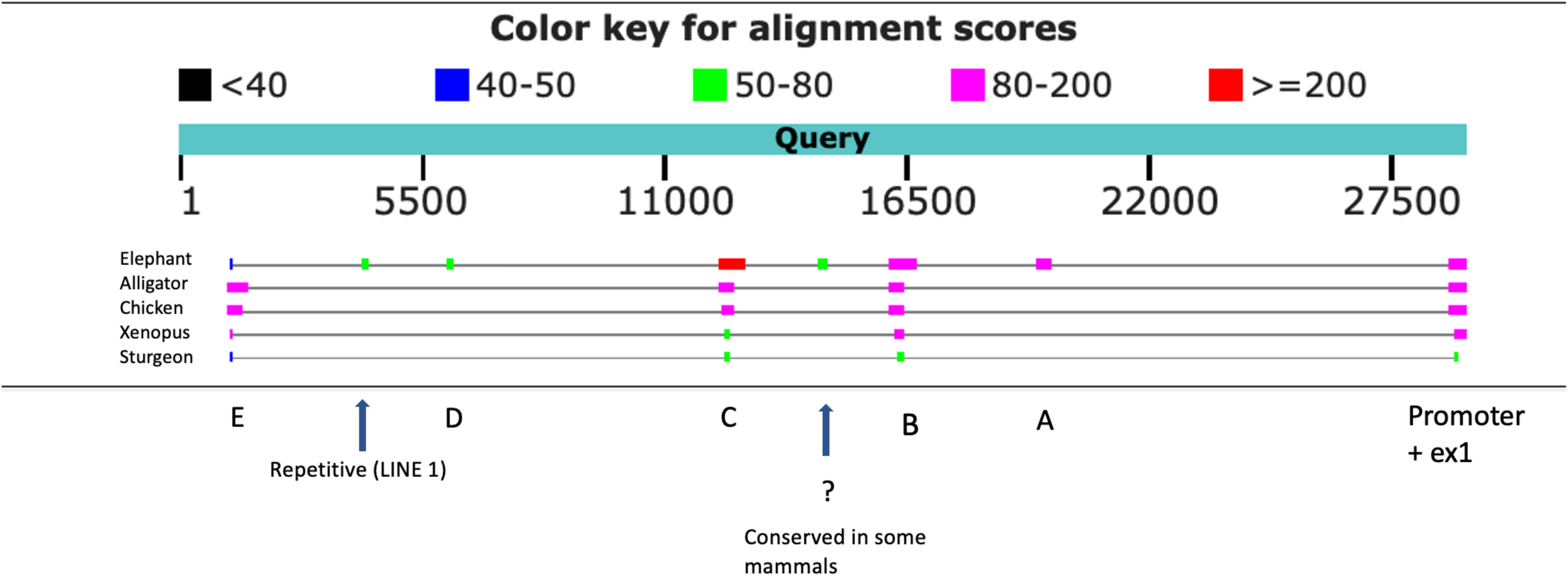
Blast2seq comparison of upstream sequences from 6 species with opossum as Query.

## Notes

### Competing Interest Statement

The authors have declared no competing interest.

